# Coordinated cerebellar climbing fiber activity signals learned sensorimotor predictions

**DOI:** 10.1101/281055

**Authors:** William Heffley, Eun Young Song, Ziye Xu, Benjamin N. Taylor, Mary Anne Hughes, Andrew McKinney, Mati Joshua, Court Hull

## Abstract

The prevailing model of cerebellar learning states that climbing fibers (CFs) are both driven by, and serve to correct, erroneous motor output. However, this model is grounded largely in studies of behaviors that utilize hardwired neural pathways to link sensory input to motor output. To test whether this model applies to more flexible learning regimes that require arbitrary sensorimotor associations, we have developed a cerebellar-dependent motor learning paradigm compatible with both mesoscale and single dendrite resolution calcium imaging in mice. Here, we find that CFs are preferentially driven by and more time-locked to correctly executed movements and other task parameters that predict reward outcome, exhibiting widespread correlated activity within parasagittal processing zones that is governed by these predictions. Together, such CF activity patterns are well-suited to drive learning by providing predictive instructional input consistent with an unsigned reinforcement learning signal that does not rely exclusively on motor errors.

## Introduction

A key role of the cerebellum is to form predictive associations between sensory inputs and motor outputs. These sensorimotor predictions are critical for generating well-timed and accurate movements, and in the absence of cerebellar function, the lack of such predictive motor output severely impairs our ability to generate coordinated responses to stimuli in the external world.

Classic models posit that the cerebellum generates sensorimotor predictions according to a supervised learning rule^1–3^. According to such models, projections from the inferior olive called climbing fibers are thought to signal motor errors, thus providing information to Purkinje cells about discrepancies between the expected consequences of a motor command and subsequent sensory feedback. To correct erroneous motor output, climbing fibers instruct heterosynaptic long-term depression^4,5^ by producing powerful regenerative calcium transients^6^ in Purkinje cell dendrites called complex spikes^7^. In so doing, climbing fibers are thought to appropriately update the cerebellar forward internal model with revised sensorimotor predictions.

This supervised error-signaling framework provides a compelling explanation for climbing fiber activity in a variety of simple behaviors, such as classical conditioning (e.g. eyeblink conditioning) and adaptation (e.g. vestibulo-ocular reflex gain changes)^8–10^ paradigms. Such behaviors typically rely on a yoked relationship between unconditioned sensory input and motor output, allowing the cerebellum to utilize signals from hardwired pathways to drive learning. Hence, the climbing fibers can instruct learning by responding to an unconditioned stimulus (e.g. periocular eye puff or retinal slip) that produces the same movement requiring modification (e.g. eyelid closure or eye movement).

However, many forms of motor learning do not involve modifications to motor programs linked directly to an unconditioned stimulus and response. Instead, the correct association between sensory input and motor output must be learned through experience, and the sensory information necessary for learning may have no direct relationship to the movement that requires modification. Such abstract associations necessitate that learning cannot be achieved by input from hardwired pathways alone. In these cases, where an unconditioned stimulus and response alone do not contain sufficient information to guide learning, it is unclear how a supervised error signal could be generated, or whether such a learning rule could account for either climbing fiber activity or the cerebellar contribution to learning.

To test how the climbing fiber system is engaged under conditions where the sensory and motor signals necessary to drive learning are not innate, we have established a cerebellar-dependent behavioral paradigm compatible with population level *in vivo* calcium imaging, optogenetic and electrophysiological approaches. Using this paradigm, we reveal two key features of climbing fiber (CF) driven complex spiking. First, we find that complex spiking cannot be accounted for by a simple error-based supervised learning model. Instead, complex spiking can signal learned, task specific predictions about the likely outcome of movement in a manner consistent with a reinforcement learning signal. Second, population level recordings reveal that while complex spiking is correlated within parasagittal zones, these correlations also depend on behavioral context. While previous measurements have shown increased correlations in complex spiking in response to sensory input or motor output^11–14^, and have suggested an important role for synchrony in downstream processing and motor learning^15^, our results reveal that such modulation can vary for identical movements depending on behavioral relevance. Hence, these data reveal key features of cerebellar CF activity that differ significantly from many classically studied cerebellar behaviors, and suggest an extension to current models of cerebellar learning in order to account for the role of complex spiking in tasks that require abstract sensorimotor associations.

## Results

To measure climbing fiber driven complex spiking, we first designed an appropriate task. An important feature of the task design is to specifically engage neurons near the dorsal surface of the cerebellum, thereby allowing for visualization of complex spikes via calcium imaging. Hence, we used optogenetics to functionally map the midlateral dorsal surface of the mouse cerebellum according to the motor output produced by spatially defined populations of Purkinje cells (PCs). To do so, we expressed Archaerhodopsin (Arch) using a transgenic approach (methods) in cerebellar PCs (Fig. 1A), and used an external fiber coupled laser to transiently suppress PCs near the dorsal surface of the cerebellum (Fig. 1B). Consistent with previous work^16^, we identified a population of superficial PCs in lobule simplex capable of driving ipsilateral forelimb movements either during optogenetic silencing or following a brief train of electrical simulation (Fig. 1, Supp. Fig. 1, Supp. Video 1). *In vivo* single unit recordings confirmed that our optogenetic stimulation was effective in silencing superficial PCs at the threshold for driving forelimb movement (~20 mW) up to a depth of 500 μm (Fig. 1B).

**Figure 1.**
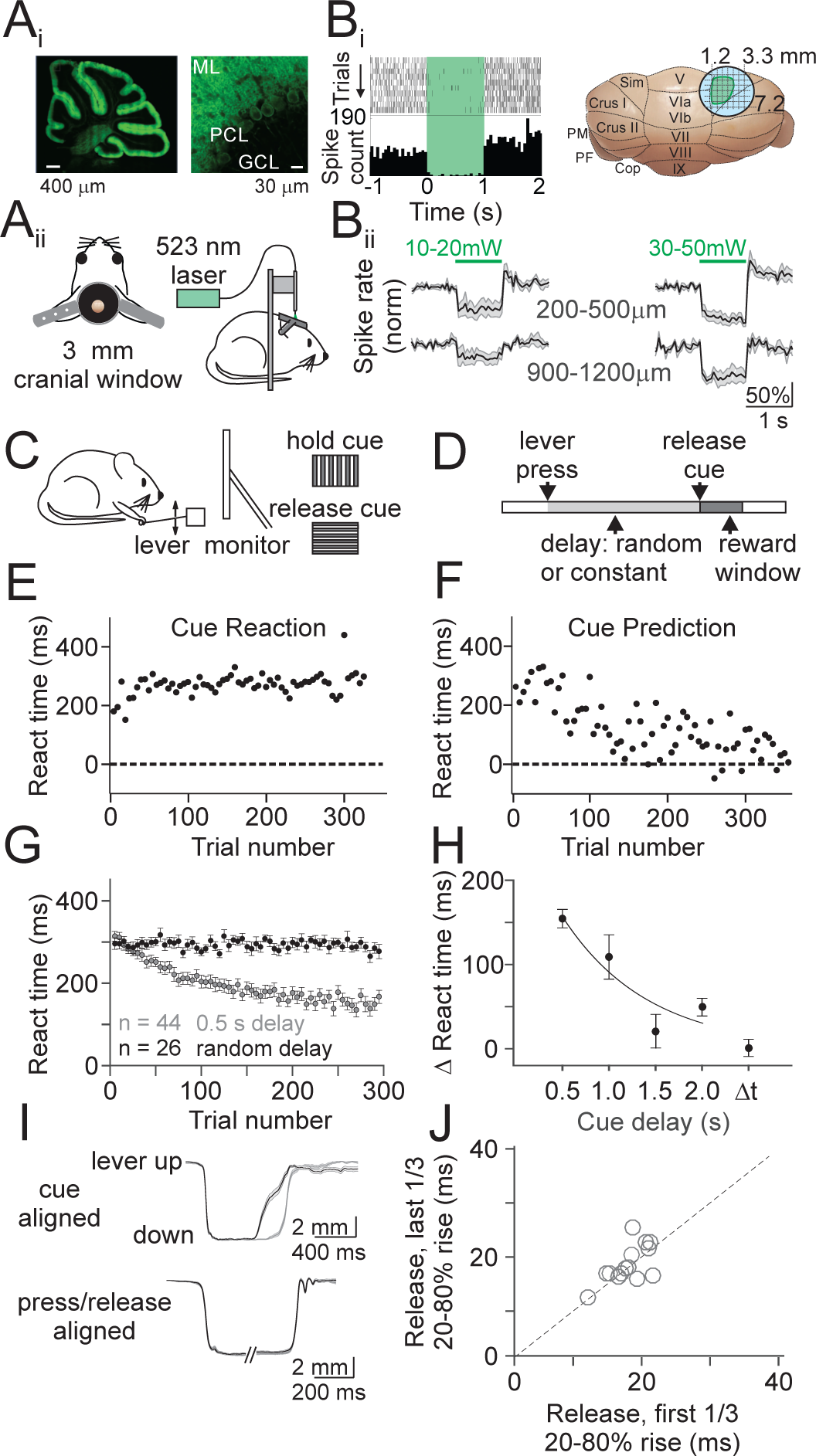
Cerebellar sensorimotor task for head-fixed mice. Ai) Confocal image of a sagittal section from PCP2-Cre x Arch mouse showing expression of the inhibitory opsin Arch in PCs at low (left) and high (right) resolution. ML; molecular layer, PCL; Purkinje cell layer, GCL; granule cell layer. Aii) Schematics of a cranial window implanted over lobule simplex (left) and configuration for optogenetic stimulation (right). Bi) *Left*, representative single unit recording of PC simple spike raster (top) and spike rate histogram (bottom) aligned to onset of Arch activation (green shaded interval). *Right*, Location of the stimulation sites that evoked ipsilateral forelimb movements (green). Bii) Average normalized firing rates from superficial (top) or deep (bottom) PCs at low (left) or high (right) stimulation powers (n=5 mice; 10-20 mW – superficial: n=6 cells; deep: n=5 cells; 30-50 mW – superficial: n=10 cells; deep: n=4 cells). Note that the lower powers used to map motor responses were only sufficient to strongly suppress neurons above 500 μm. C) Schematic of configuration of behavioral task. Head-fixed mice were trained to release a lever in response to a visual cue, with reward delivered immediately upon correctly timed movement. D) Schematic of trial structure. The delay between lever press and release cue was either randomized (cue reaction) or constant (cue prediction) from trial to trial. E) Average reaction time (from cue onset) as a function of trial number for an example cue reaction session. Each point is the average of 5 trials. F) Same as E, for a cue prediction session. G) Summary of reaction times for all cue prediction sessions with a 0.5 s cue delay (gray; n=44 sessions, 10 mice) and all cue reaction sessions (black; n=26 sessions, 7 mice). Error bars are ±SEM across sessions. H) Summary of mean change in reaction time from the beginning to the end of cue prediction and reaction sessions (methods) (n =44 sessions, 0.5 s; n=9, 1.0 s; n=6, 1.5 s; n=30, 2.0 s; n=26, Δt). One-way ANOVA, main effect of cue delay, F=10.72, df=3. Error bars are ± SEM across sessions. I) *Top*, average lever kinematic traces from an example 0.5 s cue prediction session aligned to the cue show that lever releases occur sooner in the last 1/3 of trials (black, n=149) as compared to the first 1/3 of trials (gray, n=149). Shaded area is ±SEM across trials. *Bottom,* same example session as above aligned separately to press and release showing no difference in kinematics from beginning (gray) to end (black) of the session. J) Summary of average lever release times (20-80% rise time) across sessions (n=15 sessions, 4 mice)

### A cerebellar dependent sensorimotor association learning task

We next designed a sensorimotor association task involving forelimb movement, with the rationale that the cerebellar contribution to learning in such a task should necessarily involve the PCs capable of driving forelimb movements. This task contains two discrete conditions; one where motor output is reactive (‘cue reaction’), and another where motor output becomes predictive with learning (‘cue prediction’). In both task variations, head restrained, water deprived mice are trained to self-initiate trials by depressing a lever, and to release the lever in response to a visual cue (Fig. 1C,D).

In the cue reaction condition, trial-to-trial variability in the timing of the visual cue (Fig. 1E,G, Supp. Video 2) is imposed to necessitate that mice employ reactive forelimb responses. In this regime, reaction times are reflective of the latency for sensory integration^17^ and remain constant throughout each session (Fig. 1H).

In the cue prediction condition, by presenting the visual cue with a constant delay on every trial, mice learn to predict the timing of the cue and adjust their motor responses to more closely approximate its timing (Fig. 1F,G). Moreover, learning occurs within single training sessions, is retained across days (Supp. Fig. 2A), and can be extinguished by returning mice to the cue reaction paradigm (Supp. Fig. 2C,D). In addition, we find that learning depends on the duration of the cue delay: maximal learning occurs at short cue delays (e.g. 500 ms) and little or no learning occurs for cue delays greater than 1.5 seconds (Fig. 1H; one-way ANOVA, main effect of cue delay, p = 4.8×10^−6^. Post hoc comparison to cue reaction: 500 ms, p=1.9×10^−11^, 1 s, p=8.4×10^−11^, 1.5 s, p=0.359, 2 s, p=0.005). This dependence on a short interval delay is consistent with the temporal characteristics of canonical cerebellar-dependent sensorimotor associative learning^18^.

**Figure 2.**
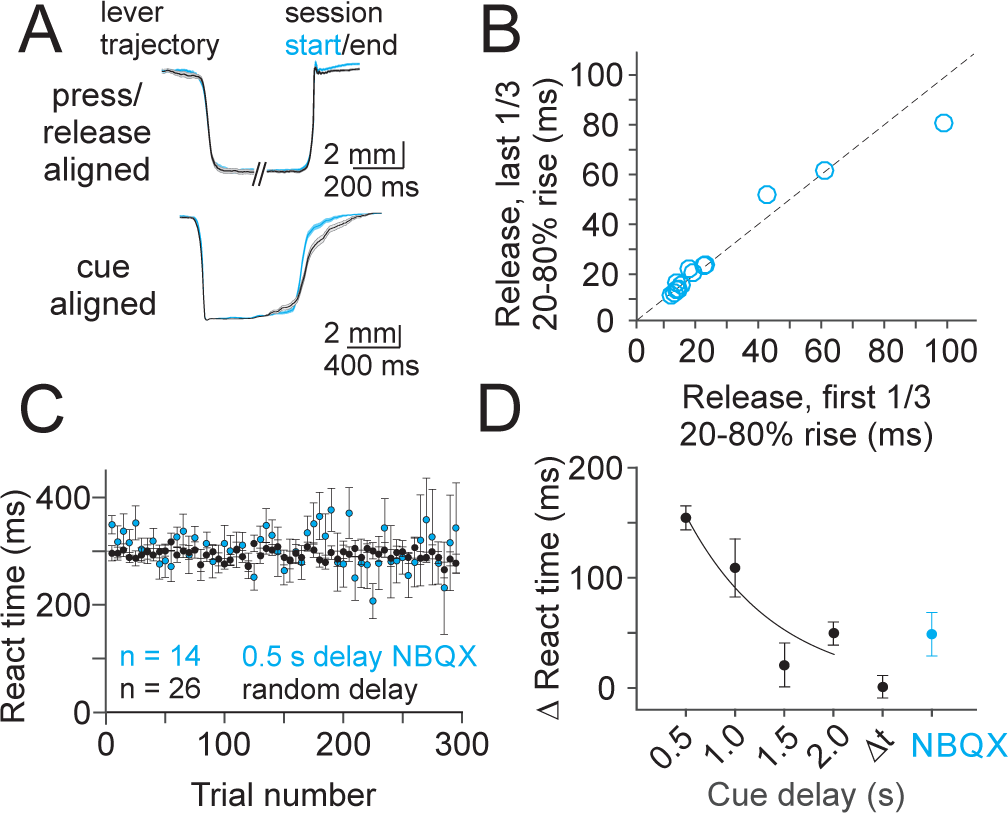
Learning requires synaptic transmission in lobule simplex. A) Average lever kinematic traces from an example 0.5 s cue prediction session after application of NBQX (300 μM), CPP (30 μM) and MCPG (30 μM). *Top*, traces aligned separately to press and release showing no difference in kinematics from beginning (blue, first 1/3 of trials, n=190) to end (black, last 1/3 of trials, n=190) of the session. *Bottom*, traces aligned to the cue show that lever releases do not occur sooner in the last 1/3 of trials (black) as compared to the first 1/3 of trials (blue). Shaded area is ±SEM across trials. B) Summary of average lever release times (20-80% rise time) across NBQX 0.5s cue prediction sessions (n=14 sessions, 6 mice). C) Summary of mean reaction time for all 0.5 s cue prediction sessions where NBQX (n=14) CPP (n=14) and MCPG (n=8) were applied locally to LS (blue; n=6 mice) compared to summary of cue reaction sessions (black; replotted from Fig 1G). Error bars are ± SEM across sessions. D) Summary of mean change in reaction time from the beginning to the end of NBQX 0.5s cue prediction sessions (blue; n=10 sessions, 6 mice) compared to cue prediction and reaction sessions (replotted from Fig. 1H). Error bars are ± SEM across sessions.

To test whether animals adjust the timing or kinematics of forelimb movement to predict the visual cue, we measured lever kinematics during a subset of learning sessions. Alignment of the lever trajectory to the cue onset confirmed that animals released the lever sooner in anticipation of the visual cue (Fig. 1I; p = 1.2×10^−5^, paired t-test). However, despite substantial variability in the kinematics of release (Supp. Fig. 3), independent alignment of both press and release demonstrated that there was no change in movement kinematics across learning sessions (Fig 1I-J; p=0.513, paired t-test).

**Figure 3.**
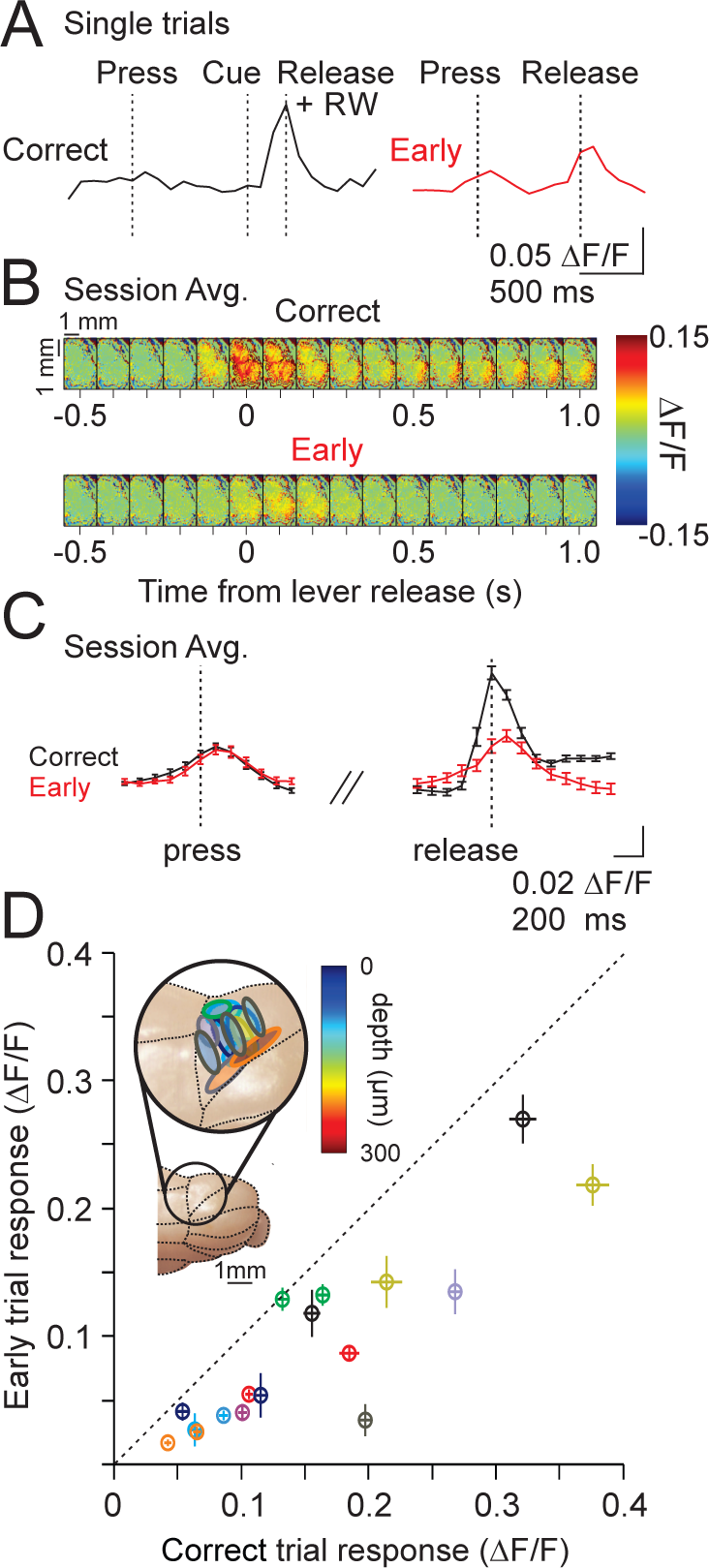
Single photon imaging during cue reaction sessions. A) Example ΔF/F single trial timecourses for a correctly timed lever release (left) and an early lever release (right). RW = reward, delivered immediately on correct release. B) Field of view (images separated by 100 ms) showing the average fractional change in fluorescence aligned to lever release for correct (top) and early (bottom) trials from an example session (same session as example from A). C) Calcium transient timecourses averaged across 87 trials from the experiment in A. B) for correct (black) and early trials (red). Error bars are ±SEM across trials. D) Summary of the mean peak calcium transients (measured on single trial basis, methods) for all correct and early trials across sessions (n=10 animals, 17 sessions). Colors represent different animals, and each point represents an imaging session. Note that all points lie below the diagonal. *Inset*, schematic of ROIs for all sessions. Outline color corresponds to mouse from scatter plot, and fill color indicates imaging depth identified by post-hoc histology. Error bars are ± SEM across sessions.

To test the necessity of lobule simplex for this learning, we first locally applied lidocaine to block spiking in cerebellar cortical neurons, including PCs. However, consistent with the critical tonic inhibition PCs provide to the deep cerebellar nuclei, we observed significant motor deficits including slowed movement, dramatically fewer initiated trials, and dystonic limb contractions (not shown). Thus, this manipulation was not appropriate to define the necessity of lobule simplex in our task.

To avoid significant motor impairment, we next used a pharmacological manipulation to selectively disrupt excitatory synaptic transmission in lobule simplex without abolishing PC pacemaking. For these experiments, we optimized concentrations of the excitatory synaptic transmission blockers NBQX (300 μM), CPP (30 μM) and MCPG (30 μM) to avoid visible impairment of motor output, and then tested this cocktail during behavior. Following local drug application (methods), mice retained the ability to release the lever in response to the visual cue, maintaining stable kinematics across sessions (Fig. 2A,B) that were not different from control sessions (Fig. 1I,J, and Supp. Fig. 3; Press p=0.261; Release p=0.222, unpaired t-test). However, despite stable movement kinematics, learning was severely impaired when the cue was presented with a constant 500 ms delay (Fig 2A,C,D) (NBQX vs Control 500 ms p=4.6×10^−4^, NBQX vs Control Cue Reaction p=0.05, unpaired t-test). Acute single unit PC recordings during drug application revealed that this manipulation significantly reduced complex spike rates, consistent with a reduction in the amplitude of postsynaptic glutamatergic currents produced by climbing fiber input (Supp. Fig. 4; p=8.5×10^−4^, paired t-test). We also found a trend toward decreased simple spiking, consistent with reducing synaptic input from the mossy fibers and parallel fibers (p=0.36, paired t-test). Hence, these data support that hypothesis that synaptic transmission in lobule simplex is necessary for predictive sensorimotor learning in our task, thus implicating cerebellar cortical learning.

**Figure 4.**
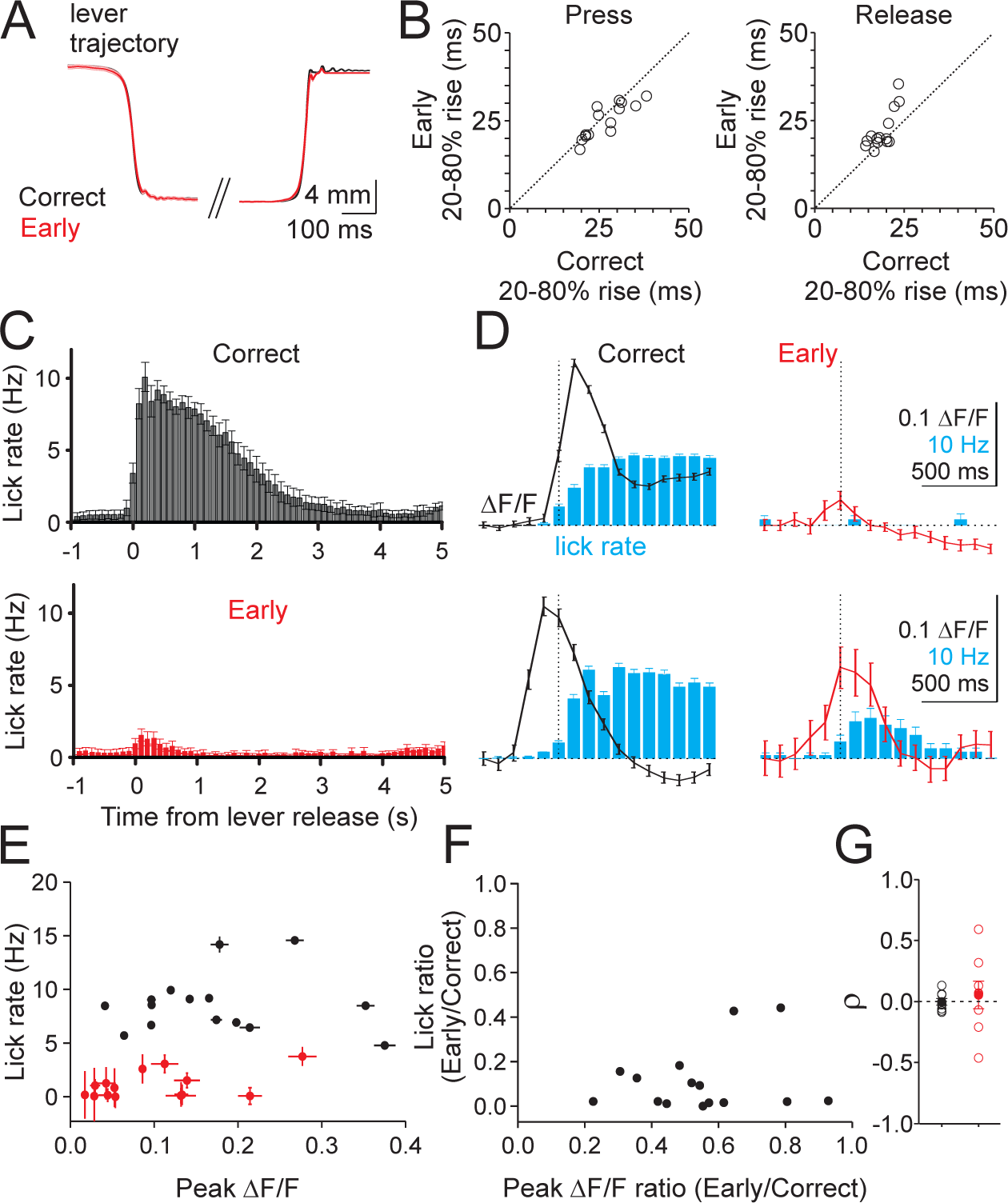
Lever dynamics and licking do not explain differences in complex spiking across trial types. A) Average lever kinematics across 385 trials from a representative cue reaction session for correct (black) and early (red) trials aligned to threshold crossing for both press and release. B) Summary of rise times for press (left) and release (right) on correct and early trials across sessions (n=14 sessions, 3 mice). C) Average timecourse of licking for correct (top) and early (bottom) release trials across sessions (n=15 sessions, 9 mice). Error bars are ±SEM across sessions. D) Two example sessions (top, n=198 trials, bottom, n=173 trials) illustrating the relationship between release-evoked calcium transients (black and red lines) and licking (blue bars) for correct (left) and early (right) release trials. Error bars are ± SEM across trials. E) Summary of the relationship between lick rates and the magnitude of fluorescent transients for correct (black) and early (red) release trials (n= 15 sessions, 9 mice). Error bars are ±SEM across trials. F) Summary of the mean ratio of early and correct trial-evoked peak calcium transient and lick rates for the same sessions in E. G) Summary of the Spearman’s correlation (ρ) between lick rate and peak ΔF/F on correct (black) and early (red) lever releases across trials for each session (n=8 sessions, 7 animals). Filled circles are mean and SEM across sessions.

### Complex spiking is not driven by motor errors in a voluntary motor learning task

Because climbing fibers provide key instructional signals for cerebellar learning, we next sought to test what information is conveyed by the olivary system in this behavior. Thus, we imaged climbing fiber evoked calcium signals in mice engaged in the cue reaction condition. In this regime, all sensorimotor signals are in place to drive learned, predictively timed forelimb movements, but no learning can occur due to the variable cue timing (Fig. 1G,H). Hence, we can test what task features drive complex spiking in a stable regime. We began by using a single photon microscope to image populations of PCs virally expressing GCaMP6f (methods). Image segmentation in a subset of experiments with the most superficial GCaMP expression revealed structures consistent in size and shape with PC dendrites in cross section, and these structures exhibited calcium transients at rates consistent with the baseline rate of complex spiking measured with single unit electrophysiology (Supp. Fig. 5). These results are consistent with several *in vivo* calcium imaging studies showing that PC responses are dominated by the large dendritic calcium transients produced by complex spikes^11-14, 19–24^.

**Figure 5.**
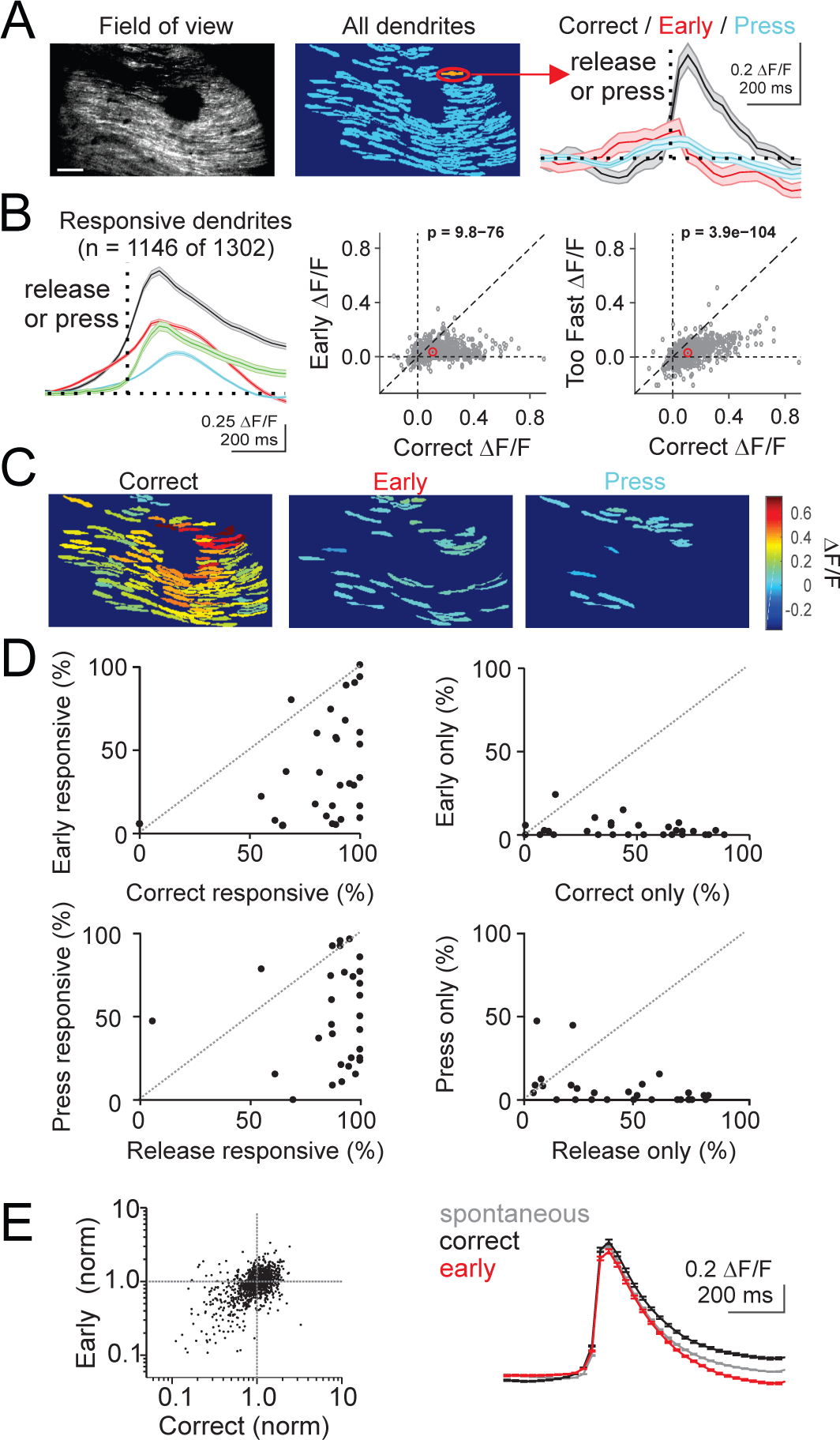
Complex spiking produces larger mean response in individual dendrites and enhanced population responses when movement is correctly timed. A) *Left*, average GCaMP fluorescence from an example 2-photon imaging session (30 Hz acquisition rate). Scale bar = 100 μm. *Middle*, pixel mask of individual PC dendrites extracted from the same session. Right, average calcium responses measured from the highlighted dendrite segregated by trial event (n=186 trials). Dotted line represents time of lever press (blue) or lever release (correct: black; early: red). Shaded area is ±SEM across trials. B) Left, average timecourse of calcium response for all significantly responsive dendrites aligned to lever press (blue), or release (correct releases (black), early releases (red), and too-fast releases (green) that occurred < 200 ms following the visual cue; n=17 animals, 30 sessions, 1146 dendrites). Shaded area is ±SEM across dendrites. *Middle*, summary scatter plot comparing the average peak amplitude of the calcium transient for each significantly responsive PC dendrite (gray) and the average of all dendrites (red) for correct and early release trials. P-value from paired t-test. *Right*, same as middle for correct and too fast release trials. P-value from paired t-test. C) Pixel masks of dendrites significantly responsive to lever release on correct trials, lever release on early trials, and lever press for the example session in A. Color map represents average ΔF/F. D) Summary across experiments of the fraction of dendrites responsive to correct lever releases vs early releases (top, left), to only correct or only early releases (top, right), to lever press vs release (bottom, left), and only to press or release (bottom, right). Responses were categorized as significant (p<0.05) according to a one-tailed t-test (methods). E) *Left,* average amplitude of correct and early release-evoked calcium events for all dendrites normalized to the amplitude of spontaneous calcium events. One-way ANOVA, F=1.8, df=2. *Right,* mean spontaneous (gray, n=286,964 events, 1146 dendrites) and correct (black, n=19,550 events, 1146 dendrites) and early (red, n=9,167 events, 1146 dendrites) release lever-evoked calcium events across all responsive dendrites. Error bars are ±SEM across dendrites.

We next identified regions of interest, defined by the expression pattern of GCaMP in each animal, that were restricted to the region within lobule simplex capable of driving ipsilateral forelimb movements (methods). Consistent with the role of this region in the control of forelimb movements, we observed task modulated PC calcium transients associated with the timing of lever press and release, with the largest modulation at the time of lever release (Fig. 3A-C). Surprisingly, when we segregated trials according to outcome, we observed significantly larger calcium transients at the time of lever releases when animals correctly released the lever in response to the visual cue (Fig. 3A-D; p=4.60×10^−5^, paired t-test). These enhanced responses were distributed widely across dorsal, superficial lobule simplex, and were consistent across sessions and animals. The same pattern of enhanced complex spiking on correctly timed lever releases was observed in the cue prediction condition, and was maintained across learning sessions, as measured by single unit PC electrophysiological recordings (Supp. Fig. 6). Thus, in this task, our data suggest that complex spikes occur preferentially on correctly timed movements, and not following motor errors.

**Figure 6.**
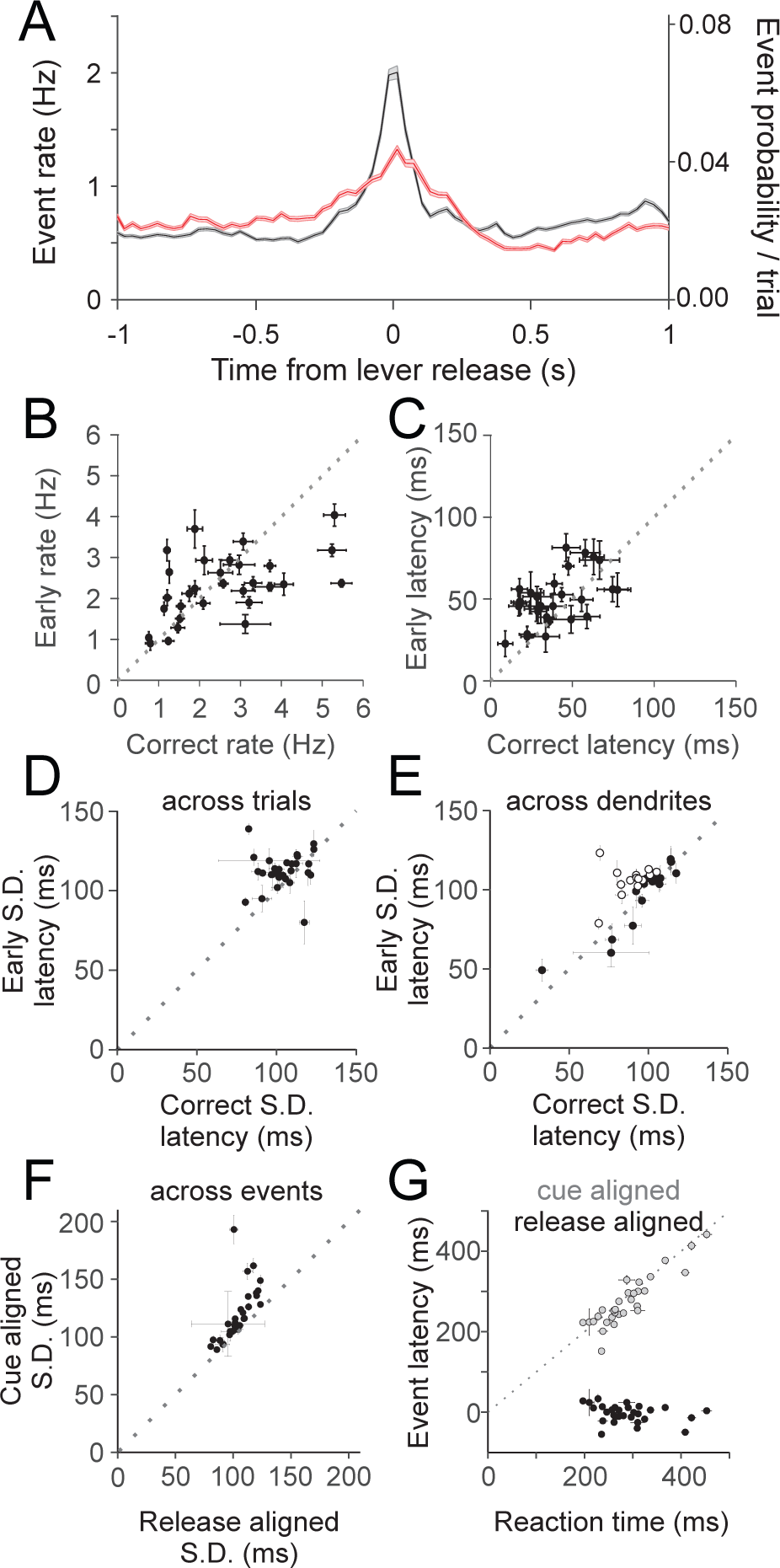
Complex spiking occurs with higher peak rates and greater synchrony when movements are correctly timed. A) Peri-release time histogram of calcium events on correct (black) and early (red) release trials (n=17 animals, 30 sessions, 1146 dendrites). Bin width = 33 ms. Shaded area is ±SEM across dendrites. B) Summary of average event rate for correct and early releases for each session (n=30). Error bars are ±SEM across dendrites. C) Same as B for average event latency relative to release for each session (n=30). Error bars are ±SEM across dendrites. D) Summary of average standard deviation (S.D.) of event times across trials within single dendrites for each session (n=30). Error bars are ±SEM across dendrites. E) Same as D for measurement of S.D. of event times across dendrites for each session (n=30). Open circles are statistically significant for reduced correct trial jitter across dendrites (paired t-test). Error bars are ±SEM across trials. F) Summary of the average S.D. of event times when aligned to either lever release or visual cue for each session (n=30). Error bars are ±SEM across dendrites. G) Summary of the average latency to peak event probability relative to cue (gray circles) or lever release (black circles) compared with the average session reaction time for each session (n=30). X-error bars are ±SEM across trials; Y-error bars are ±SEM across dendrites.

There are many differences between correct and early release trials that could contribute to the outcome-dependent difference in complex spiking. One possible difference between trial types could be the kinematics of forelimb movement. Hence, in a subset of our experiments, we measured movement kinematics by comparing lever trajectories on correct and early release trials (Fig. 4A-B). These data revealed closely matched movement kinematics between correct and early lever release trials (Press p = 0.06, Release p = 0.007, paired t-test).

Another difference between correctly timed and early lever releases is that reward is only delivered on correct trials, and this results in extended licking. Indeed, we find that lick rates were significantly higher on correct trials (Fig. 4C; p=3.74×10^−9^, paired t-test). Moreover, in some sessions we observed a late, sustained calcium transient that occurred during licking (Fig. 4D). However, we found no correlation between the mean peak calcium transients and mean lick rates across trial types (Fig. 4E,F; Correct p=0.11; Early p=0.98, linear regression) or within sessions across trials of each type (Fig. 4G; Correct p=0.93; Early p=0.65, one sample t-test; Supp. Fig. 7D; Correct p=0.544; Early p=0.428, one sample t-test). Thus, we conclude that the enhanced responses as measured by the peak calcium transient on correctly timed lever releases are not due to differences in forelimb movement kinematics or licking (Fig. 4 and Supp. Fig. 7), but are instead linked to the context of the movement.

**Figure 7.**
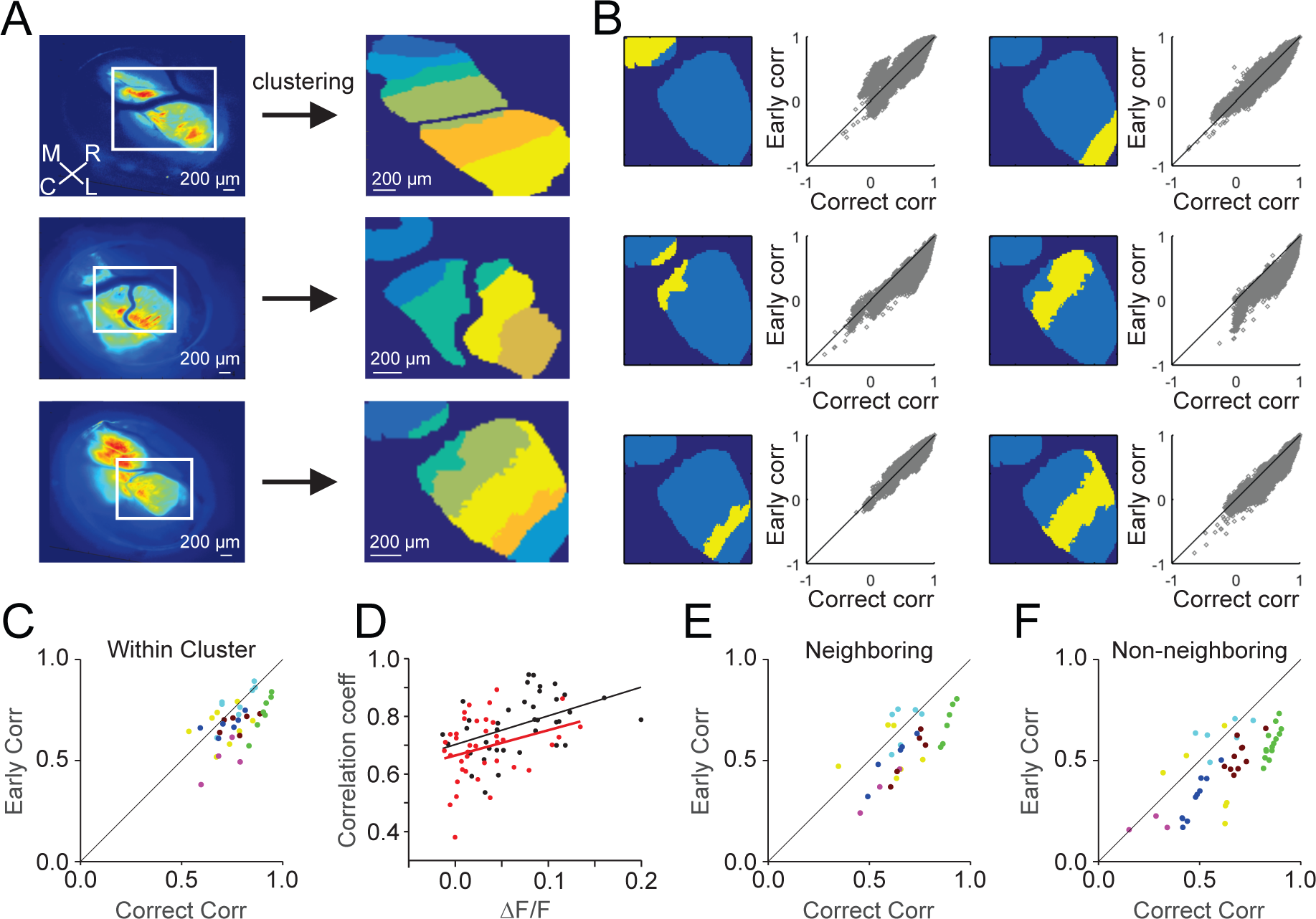
Complex spiking is correlated across parasagittal zones, with higher correlations on correct lever releases. A) *Left*, Average GCaMP fluorescence for three example experiments. ROIs from lobule simplex (white boxes) were selected for unsupervised clustering analysis (meta K-means, methods). *Right*, pixels are colored according to cluster identity for each example experiment. M=medial, R=rostral, C=caudal, L=lateral. Note the alignment of the clustered zones along the rostro-caudal axis. B) Within cluster correlations (n=6 clusters, Pearson’s) between correct and early lever release trials from an example session in A (bottom) for each of the identified clusters (yellow). C) Summary of average Pearson’s correlation coefficient for all pixel pairs (methods) within clusters (n = 39) across experiments for correct and early release trials (n=5 animals, 7 sessions, 39 clusters). Colors denote clusters from the same session. D) Summary of the relationship between peak ΔF/F and Pearson’s correlation coefficients for paired correct (black) and early (red) trials for each cluster (n=39). Linear fits were performed for within cluster correlations using correct and early trials separately. E) Same as C for pairs of pixels across neighboring clusters (n=31 cluster pairs). F) Same as C for pairs of pixels across non-neighboring clusters (n=50 cluster pairs).

### Behavioral context determines the probability and synchrony of complex spiking

To investigate the dynamics of complex spiking at the single cell level, we used video-rate two-photon microscopy to image the dendrites of PCs during behavior. Using this approach, we isolated the dendrites of individual PCs^20^ (Fig. 5, Supp. Figs. 5, 8), and measured both spontaneous and task-evoked calcium transients. As with the single photon imaging experiments, spontaneous calcium transients measured during inter-trial intervals occurred at a rate of 0.61 ± 0.01 Hz, consistent with our electrophysiological measurements of spontaneous complex spiking (Supp. Fig. 5E,F). The two-photon data also replicated the main finding that the mean calcium transients were significantly larger on correct release trials within individual PC dendrites (Fig. 5A,B; p=9.79^−76^, paired t-test). Moreover, lever releases that were rewarded but occurred too fast to be responses to the visual cue (reaction times < 200 ms) produced calcium transients that were significantly smaller than those produced by correct lever release trials (p=3.91^−104^, paired t-test) and not significantly different from responses on early release trials (p=0.086). Hence, these data support the analysis in Figure 4 that the enhanced complex spiking on correct trials is not due to licking or reward per se.

**Figure 8.**
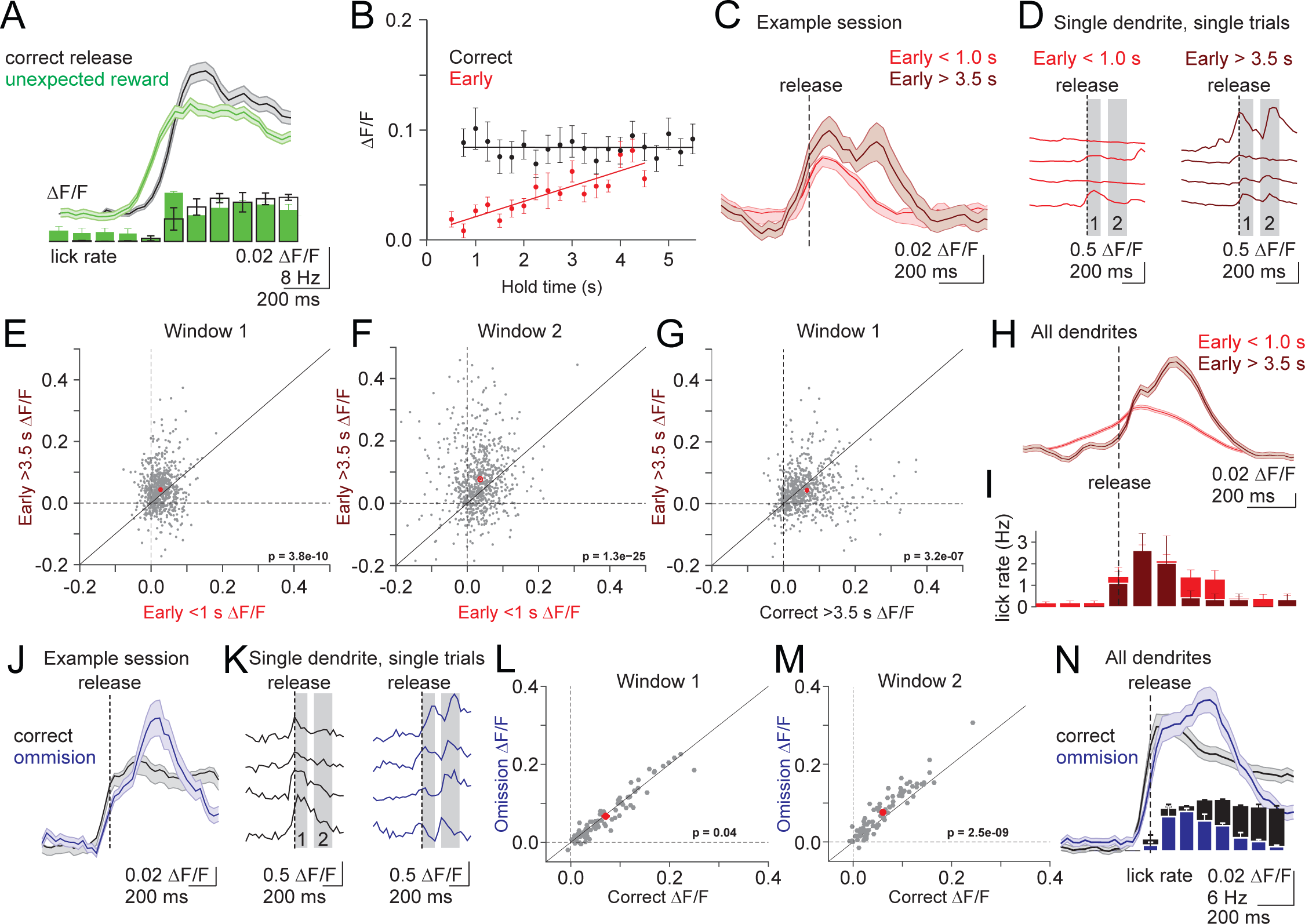
Complex spiking is modulated by learned sensorimotor predictions. A) Top, Average calcium transient in response to unexpected reward (green, aligned to first lick) and correct lever releases (black, aligned to release). Shaded error is SEM across dendrites (n=120). Bottom, Average lick rate for unexpected reward and correct lever releases. Error is SEM across sessions (n=3). B) Summary of the mean peak calcium transients in a 500 ms window at the time of release (methods) across all 2-photon experiments for correct (black) and early (red) release trials binned according to hold time (250ms bins). Linear fits were applied to data from each trial type (n=17 animals, 30 sessions). Error bars are SEM across sessions. C) Mean timecourse of calcium transients for an example session (n=43 dendrites) divided according to early trials with hold durations of < 1.0s (red) and > 3.5s (dark red). Note the late, second response for long hold duration trials. Shaded error is SEM across dendrites. D) Single trial example timecourses from an individual dendrite from the session in C. Gray bars represent analysis windows for subsequent plots E-G) Summary of peak calcium transients measured in the shaded windows from D for all dendrites (n=1146) according to lever hold time (E,F) and trial outcome (G). P-values represent paired t-tests. H) Mean calcium transient timecourses averaged across all dendrites (n=1146) on short and long hold early release trials. Shaded error is SEM across dendrites. I) Mean lick rates for the subset of experiments where licking was measured in H (n= 6 animals, 10 sessions)). Error bars are SEM across sessions. J) Example timecourses of mean calcium transients from a single session (n=22 dendrites) for correct (black), and reward omission (dark blue) trials. Note the late, second response on omission trials. Shaded error is SEM across dendrites. K) Single trial example timecourses from an individual dendrite from the session in J. L-M) Summary of peak calcium transients measured in the shaded windows in K on correct and reward omission lever trials (n=81 dendrites, 3 animals, 3 sessions). P-values represent paired t-tests. N) Mean timecourses of calcium transients averaged across all dendrites (n=81) for correct (black) and reward omission (dark blue) trials. Vertical bars represent the lick rate averaged across trials for all reward omission sessions. Shaded error is SEM across dendrites. Error bars are SEM across sessions.

In addition to larger mean responses across trials, population level analysis revealed that more PCs were driven on correct release trials (Fig 5C). Of the total PC population that responded to any lever press or release (1146 of 1302), most (88%) PCs exhibited complex spikes on correct release trials, whereas less than half (41%) responded during early lever releases (Fig. 5D, *top*). Approximately half (48%) exhibited complex spikes in response to lever press (Fig. 5D, *bottom*). The increased number of active dendrites on correct trials is unlikely to be a thresholding artifact, since there was no significant difference in the average response amplitude of correct-only and correct+early responsive dendrites to correct trials (p=0.44, unpaired t-test, correct-only n=578 dendrites, correct+early n=457 dendrites). These data also indicate that the measured increase in number of active dendrites is not due to contamination of fluorescent signals from neighboring dendrites. Hence, these single dendrite measurements explain our single photon results by demonstrating that both more PCs exhibit complex spikes on correct trials, and those dendrites that exhibit complex spikes in response to both trial types have larger mean calcium transients on correct trials.

The difference in the average release-evoked response on correct versus early trials may be due to a difference in the amplitude of the calcium transients on each trial. For instance, complex spikes are not “all-or-none” events^25^, and the number of spikelets can vary from event to event^26^, altering the size of the evoked calcium transient^27^. Thus, we extracted single trial events associated with lever releases across trial types, and compared these with each other and to spontaneous calcium transients. However, this analysis revealed no significant difference between the amplitude of calcium transients on single trials in response to either correct or early lever releases, and between movement-evoked and spontaneous activity (Fig. 5E; p=0.17, one-way ANOVA). Thus, if there are variable numbers of evoked spikelets across trials, such differences are not systematically correlated with either trial outcome or task engagement. Moreover, the equivalent amplitude of lever-evoked and spontaneous events suggests that the majority of the signal measured in response to lever release is due to complex spiking. Indeed, the size and shape of our segmented ROIs is consistent with several studies showing that complex spikes produce a calcium signal throughout the entire PC dendrite^28, 29^, whereas parallel fiber calcium signals are much smaller and confined to isolated regions of spiny dendritic branchlets^30, 31^.

To assess how equivalent single event responses can produce larger mean responses on correct trials, we generated a peristimulus time histogram of complex spike events for each trial type aligned to the time of release (Fig. 6A). This analysis revealed a significantly enhanced complex spike rate at the time of lever release when movement was correctly timed (Fig. 6A,B; p=8.14×10^−15^, paired t-test). In addition, complex spiking on correct trials in single PC dendrites occurred with shorter latency (Fig 6C; p=2.65×10^−11^, paired t-test) and less temporal jitter across trials (Fig 6D; p=0.007, paired t-test). Notably, these measures of jitter are likely to be an overestimate, since our measure of event time is limited by our sampling interval (33 ms). However, when accounting for the overall spontaneous event rate of ~1 Hz, the 100 ms jitter of events on correct trials is comparatively precise.

The PSTH reveals that elevated complex spiking occurs proximal to lever release. To test whether the increase in complex spiking is more closely associated with the lever release or the visual cue, we compared the temporal jitter of spike times on correct trials when aligned to each of these events. This analysis revealed a significantly lower temporal jitter when the spikes were aligned to lever release as measured either with imaging (Fig 6F; p=6.2×10^−5^, paired t-test) or at higher temporal resolution with electrophysiology (Supp. Fig. 6F; p=7.8×10^−9^; paired t-test). Notably, the closer association between spiking and the lever release was maintained across learning in cue prediction sessions, even as lever releases moved closer to the timing of the visual cue (Supp. Fig. 6E-F). In addition, we find that the latency to peak event rate is strongly correlated with reaction time when the events are aligned to visual cue (Fig 6G; p=8.56×10^−12^, Pearson’s correlation), but not when aligned to lever release (p=0.13). These results further suggest a closer association between lever release and complex spiking than for the visual cue and complex spiking.

The narrower PSTH on correct release trials is also consistent with the possibility of enhanced synchrony on the time scale of ~100 milliseconds at the population level. Indeed, we find enhanced population synchrony across dendrites (Fig 6E; p=0.003, paired t-test). However, in many experiments, we observed no such increase in population synchrony on correct trials (16 out of 30 sessions do not have significantly elevated synchrony on correct trials, open circles, p>0.05 for each session). We thus considered the possibility that enhanced synchrony was location specific, and might obey spatial structure that was not well demarcated at the scale of the field of view in our two-photon experiments.

### Context determines correlated population activity at the mesoscale level

To investigate whether correlated complex spiking was spatially organized, we used an unsupervised, iterative pixel-clustering approach^22^ (methods) to assess correlations between pixels across all trials at the mesoscale level from single photon imaging sessions with widespread GCaMP expression (Fig. 7A). Despite non-uniform, unpatterned GCaMP expression, this analysis revealed spatial patterns of correlated activity that were organized across parasagittal bands oriented in the rostro-caudal axis. These bands were 221 ± 15 μm wide on average across 39 measured zones, consistent with previous anatomical and physiological measurements of cerebellar microzones^32^. Hence, these data support the longstanding hypothesis that the anatomical pattern of climbing fiber projections into the cerebellar cortex establishes functionally distinct parasagittal processing modules.

To test how complex spiking is modulated during behavior within and across parasagittal zones, we divided trials according to outcome and analyzed brief epochs surrounding the time of lever release (Fig. 7B). While this analysis lacks the fine temporal resolution of the two-photon event based analysis in Figure 6, it nonetheless revealed that activity amongst pixels within a zone exhibited higher correlations for most zones on correct lever release trials (Fig. 7C; p = 1.60×10^−5^, paired t-test). These enhanced correlations are not an artifact of elevated complex spike rates on correct trials because early release trials still had weaker correlations across equivalent activity levels as assessed by linear fits to bootstrapped distributions paired from each trial type (methods) (Fig. 7D). We also found enhanced correlations between neighboring and non-neighboring zones on correct lever release trials (Fig. 7E,F; Neighboring p=0.004, Non-Neighboring p=6.2×10^−12^, paired t-test). However, these cross-zone correlations were significantly lower on average than those within zones (Within Zone vs Neighboring, p=0.004; Within Zone vs. Non-Neighboring p=3.4×10^−4^, unpaired t-test). Hence, these data reveal a precise spatial organization of climbing fiber activity, with highly correlated complex spiking within parasagittal zones that is task specific, differing for movements with the same kinematics depending on behavioral context.

### Complex spiking signals learned sensorimotor predictions

Our data suggest that complex spiking does not signal motor errors in this behavioral paradigm. Instead, our results are consistent with the possibility that the climbing fibers either 1) instruct a different type of supervised learning rule based on correctly timed motor output, or 2) provide a reinforcement learning signal, such as a temporal-difference (TD) signal of the type recently identified for climbing fibers during conditioned eyeblink learning^33^. While the instructional signals for supervised learning encode specific outcomes, those for reinforcement learning are driven by prediction errors. Thus, while a reinforcement learning signal should occur to any unexpected outcome (e.g. unexpected reward) or event that predicts task outcome, a supervised signal based on correct movements should only occur for correctly timed arm movements. We hence began by testing whether climbing fibers can be driven by unexpected reward.

Surprisingly, when reward was delivered unexpectedly during the intertrial interval, we observed robust climbing fiber responses of similar amplitude to those driven by correctly timed lever releases (Figure 8A, Supp. Fig. 9). Because we have already demonstrated that the increase in climbing fiber activity is not driven by licking or reward (Figs. 4, 5B, Supp. Figs. 7,9), we interpret the increase in complex spiking in response to an unexpected reward as evidence of a prediction error. We hence conclude that climbing fibers do not specifically represent correctly timed movement per se, but may instead provide input consistent with a reinforcement learning rule.

To further test whether climbing fiber activity in our task is consistent with a reinforcement learning signal, we next tested whether the climbing fibers respond to other task features that predict task outcome (reward delivery). Thus, we took advantage of the temporal structure of the task, wherein the probability of cue appearance, and thus reward delivery, increases with lever hold time (i.e. the hazard function is not flat). We specifically tested whether complex spiking depends on lever hold time for correct and early lever releases.

We find that while complex spiking is independent of lever hold time for correctly timed movements (Fig. 8B; 2-photon p=0.98, Supp. Fig.10; 1-photon p=0.76, Spearman’s correlation), there is a strong positive relationship between hold time and complex spiking for early releases across both single and multiphoton imaging sessions (2-photon p=4.58×10^−6^, 1- photon p<1.0×10^−16^, Spearman’s correlation). The elevated complex spiking associated with longer hold times (Fig. 8C) is due to both an increase in spiking at the time of release (Fig. 8E; Window 1 p=3.84×10^−10^, paired t-test), and an additional late response not present on short duration early release trials (Fig. 8F; Window 2 p=1.33×10^−25^; paired t-test). The late response occurred on single trials and within individual dendrites, indicating that it was not generated by a diversity of response timings across trials or by a separate population of PC dendrites (Fig. 8D). In addition, the late response occurred approximately 200 ms after lever release when reward was no longer possible and lick rates began to decrease (Fig 8 H,I). This provides additional evidence that increases in licking cannot explain the observed increases in complex spiking. Instead, this result reveals that licking is tightly linked to the animals’ expectation: when the animal no longer expects reward, lick rates decrease. In the same manner, because spontaneous licks can reflect an expectation of possible reward, it is not surprising that spontaneous licks are sometimes associated with increases in complex spiking (Supp. Fig. 7).

Each of these results are consistent with a reinforcement learning rule based on prediction error, where complex spiking increases in response to learned task events that predict reward and unexpected outcomes (i.e., the lack of reward following long duration early releases). In this model, an initial prediction accompanies lever release that depends primarily on cue presentation. Thus, complex spiking in response to correct releases is significantly stronger than on early releases, even for long duration holds (Fig. 8G; Window 1 p = 3.16×10^−7^, paired t-test). However, while the visually driven lever release provides the strongest prediction of trial outcome, trial duration also contributes to expectation according to the task structure. In the case that initial expectation is unmet, the climbing fibers also signal this unexpected outcome through an increase in spiking. Notably, this latter result argues that climbing fibers encode an unsigned prediction error, in which there is an increase in spiking regardless of the direction of the prediction error.

To further test this model, we performed another set of experiments to probe the role of violated expectations in driving complex spiking. In these experiments, we omitted reward delivery on 20% of correctly timed lever releases (Fig. 8 J-N). Consistent with the cue appearance establishing expectation at the time of lever release, both rewarded and omission trials resulted in calcium transients that were nearly equivalent at the time of lever release (Fig. 8L; Window 1 p=0.04, paired t-test). However, omission trials resulted in an additional, late response at the time when reward delivery was no longer possible and lick rates began to decrease (Fig. 8M; Window 2 p=2.54×10^−9^, paired t-test). The timing of this late response was similar to the late response following long duration early releases, and was present on single trials and within individual dendrites (Figure 8J,N). Hence, these experiments support the hypothesis that climbing fibers can both signal and evaluate predictions about the likely outcome of movements in a manner consistent with a reinforcement learning rule.

## Discussion

We have established a sensorimotor association task that involves PCs near the dorsal surface of lobule simplex. This behavior has several hallmarks of cerebellar learning, including a dependence on short delay intervals for generating the learned sensorimotor association and the requirement for excitatory synaptic transmission in the cerebellar cortex. In this behavior, however, climbing fibers do not signal erroneous motor output as described by classical models of supervised learning. Specifically, we demonstrate that climbing fiber driven complex spiking occurs with higher probability, shorter latency, and less jitter when movement is correctly timed.

Our data support the hypothesis that these enhanced climbing fiber responses are related to the predicted outcome of movement, which in this behavior constitutes delivery of a water reward. Evidence supporting this model is threefold: First, the highest probability of complex spiking occurs on correctly timed lever releases when the visual cue instructs behavior and thus a high reward expectation. Second, complex spiking is modulated by the temporal features of the task that dictate expectation. Specifically, complex spiking in response to lever release increases with increasing hold duration, matching the increased likelihood of reward as trial length increases. Third, complex spiking is also driven when the expectation of reward is violated: when the reward is omitted on correctly executed movements, not provided on longer hold duration early releases when a correct outcome of lever release is probable, or unexpectedly presented during the inter-trial interval. Together, these experiments suggest that climbing fibers carry instructional signals that both predict and evaluate the expected outcome of movement in a manner consistent with a reinforcement learning rule. Such responses are also consistent with the known role of cerebellar circuits in generating predictive motor output, and we suggest that they could thus provide a substrate for generating and testing the type forward models that have long been hypothesized to govern cerebellar processing.

The climbing fiber activity observed in response to violated expectations has some similarities to the motor error signals seen in classic cerebellar behaviors. However, in our behavioral paradigm, climbing fibers do not signal movement errors, and correct movements have the same kinematics as mistimed movements. Thus, these results indicate that climbing fibers can respond differentially to the same movement according to its expected outcome. These context-specific climbing fiber responses may be due to aspects of our task design. The behavior described here differs from most cerebellar dependent learning regimes in that the movement requiring modification (lever release) is not directly related to an unconditioned stimulus (reward) or response (licking). As a consequence, the cerebellum cannot harness sensorimotor input from hardwired pathways to enable learned changes in motor output. Instead, the necessary forward model must define and evaluate the relationship between a neutral visual stimulus, a forelimb movement, and reward.

It remains unclear whether the climbing fiber activity that enables such a forward model is generated at the level of the olive or inherited from upstream brain regions such as the neocortex, colliculus, or elsewhere. However, evidence suggests that the olive may have access to different information depending on task requirements. Anatomical and physiological work in the rodent has demonstrated that the pathway from the forelimb region of motor cortex to the olive is independent of the pathway providing ascending sensorimotor input from the periphery^34^. Such data argue that the olive has access to unique information in tasks that involve a descending motor command, and further suggest that the olive may have access to diverse cortical computations. Indeed, the presence of abstract task timing information suggests that the olive has access to higher-order signals. In support of this view, evidence from a different forelimb task in non-human primates that also requires an abstract sensorimotor association has shown similar climbing fiber responses^35^. Specifically, this study demonstrated that complex spikes can signal both the predicted destination of reaching as well as deviations from the expected destination.

Importantly, the climbing fiber activity patterns described here are appropriate to drive motor learning under conditions that require flexible sensorimotor associations. Specifically, higher probability complex spiking and enhanced correlations at the population level are both context dependent and spatio-temporally organized. At the population level, our single photon imaging has revealed correlated complex spiking within parasagittal bands of approximately 200 μm, likely corresponding to “microzones”^32^. Because PCs across microzones can respond to different sensorimotor input, and have different spiking^36^ and synaptic^37^ properties, these zones are thought to constitute discrete computational units. Recent measurements have revealed that complex spiking in microzones can become more synchronous during both motor output^14^ and sensory input^11, 12, 23^. Our population imaging results provide an extension to this idea by demonstrating that correlated complex spiking within parasagittal zones is not simply enhanced by sensorimotor input, but rather can be enhanced for the same action according to behavioral context. Precisely timed spiking across a population of nearby PCs that receive common parallel fiber input and converge to the same DCN neurons would provide an ideal substrate to maximize the impact of plasticity. However, other circuits could also play a role in learning. In particular, complex spiking can robustly inhibit nuclear neurons^38,39^, and enhanced climbing fiber synchrony would magnify this effect via convergence of simultaneously active PC axons in the DCN^40^. Because synchronous inhibition has been shown to play a key role in synaptic plasticity at nuclear neuron synapses^41,42^, correlated complex spiking across populations of nearby PCs could place the DCN as the central site of learning under such conditions^43^. In either case, we note that the climbing fiber responses following correct lever releases are well-timed to promote learning in the cue prediction condition by instructing movements that more closely approximate earliest time of reward delivery.

Finally, it is notable that ours are not the first results to suggest alternate learning rules instructed by climbing fibers. Recently, conditioned eyeblink learning data has pointed toward climbing fibers implementing a temporal difference (TD) reinforcement learning rule by signaling sensory prediction errors^33^. Because we do not observe evidence of negative prediction errors on reward omission trials or long duration early release trials, our results are incompatible with this model as strictly interpreted. Instead, our data show some similarities with the type of unsigned prediction signals observed in serotonergic^44^ and other neurons^45, 46^ that have been also thought to enable associations between unexpected outcomes and novel cues^47^. Evidence for both models can often be found in neural activity within the same brain region, and it has been suggested that TD and unsigned prediction mechanisms may in fact be linked, and act together to promote learning^48^. Hence, as has been done in other brain regions, it will be necessary to systematically vary task parameters such as the valence of instructional stimuli and the requirements for learned associations in order to resolve how the cerebellum implements discrete learning rules according to task demands beyond their classic role in supervised learning^49^. For the present, the results described here demonstrate that complex spiking can signal learned, task specific information necessary for flexible control of complex motor behaviors in a manner that does not depend exclusively on motor errors.

## Data and Code Availability Statement

Data analysis code is available from the corresponding author upon reasonable request. The data that support the findings of this study are also available from the corresponding author upon reasonable request.

## Acknowledgements

This work was supported by grants from the NIH NINDS (5R01NS096289-02) (CH) and (F31NS103425) (WH), the Sloan Foundation (CH), and the Whitehall Foundation (CH). We thank L. Glickfeld for helpful discussions and input on calcium imaging approaches and analyses, S. Lisberger, G. Field, K. Franks and F. Wang for feedback on early versions of this manuscript, and members of the Hull and Glickfeld labs for input and technical assistance throughout the project.

## Author Contributions

WH, ES, and CH designed experiments, WH, ES, AM, BT, MAH, and MJ conducted experiments, WH, ES, ZX, BT, AM, MJ, and CH analyzed data, and CH wrote the manuscript.

---

**Figure 1.**
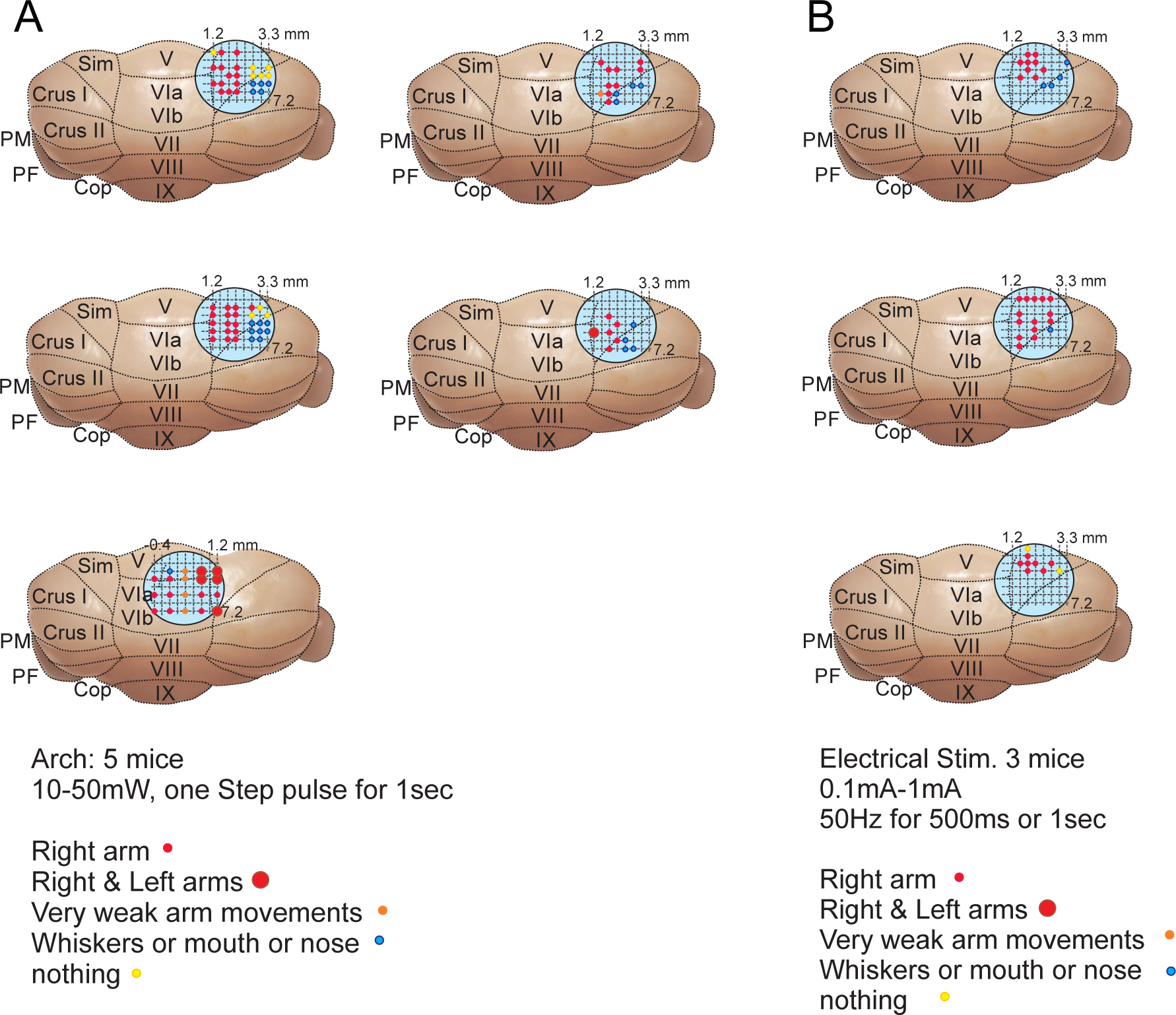
Mapping of motor output driven by Arch and electrical stimulation of superficial cerebellar cortex. A) Schematics of the results of Arch stimulation over a grid of sites in lobule simplex (Sim) and Crus I for all five mice. Colors denote outcome of stimulation. B) Same as A for three mice tested with electrical microstimulation.

**Figure 2.**
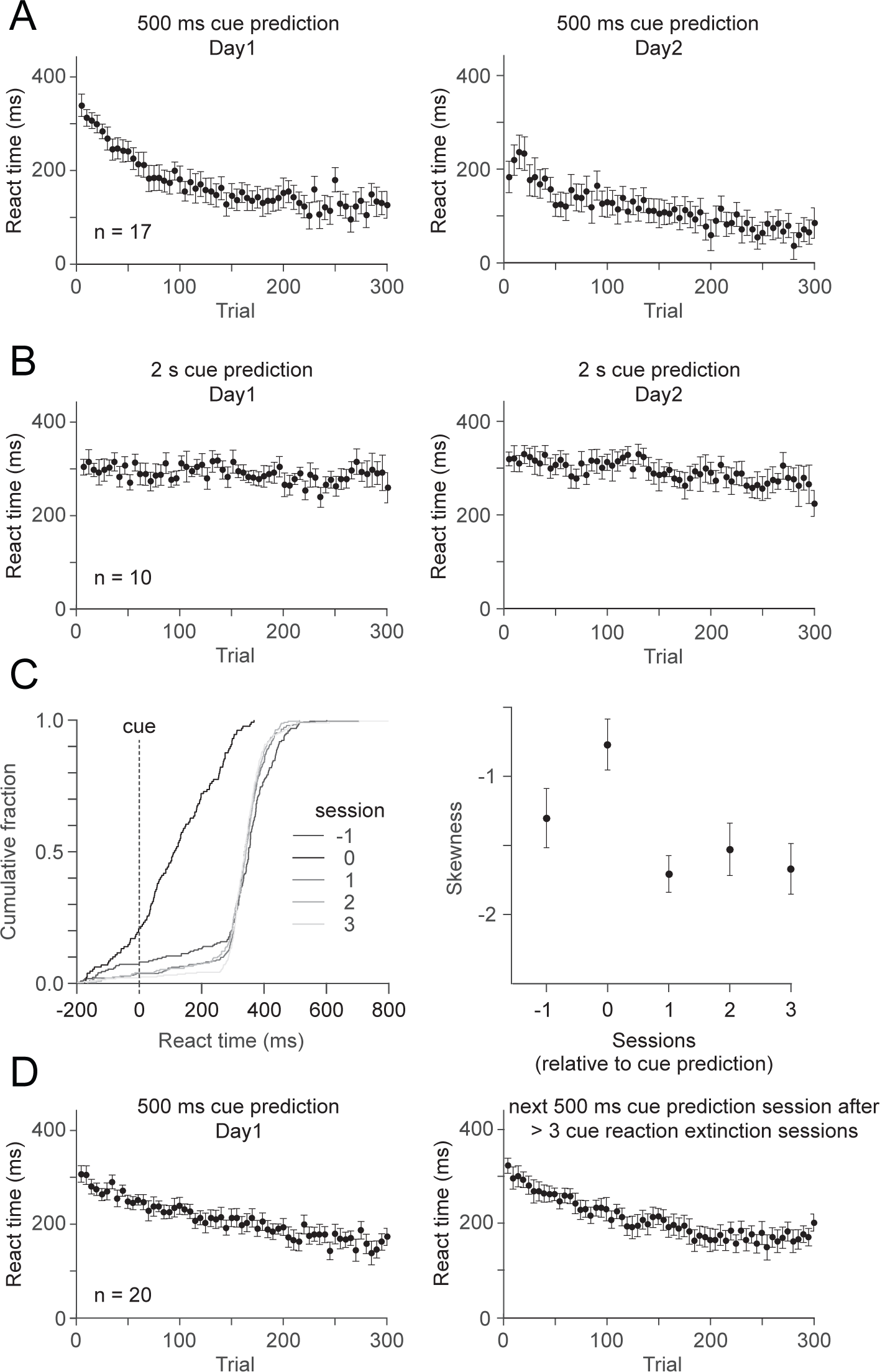
Retention and extinguishment of learning for cue prediction regime. A) Average reaction times relative to cue for the first (left) and second (right) sessions where animals performed 0.5 s cue prediction on consecutive training days (n=17 sessions, 5 mice). Error bars are ±SEM across sessions. B) Same as A for 2 s cue prediction sessions (n=10 sessions, 6 mice). C) *Left,* example cumulative distributions of reaction times for cue reaction sessions immediately surrounding a cue prediction session (session 0). Trials from the cue prediction session include only the last 1/3 of the session when the animal was anticipating the cue timing. *Right,* summary of distribution skewness across cue reaction sessions immediately surrounding cue prediction sessions (n=43 sessions). Error bars are SEM across sessions. D) Same as A for pairs of 0.5 s prediction sessions separated by 3 or more cue reaction sessions (n=20 sessions, 10 mice).

**Figure 3.**
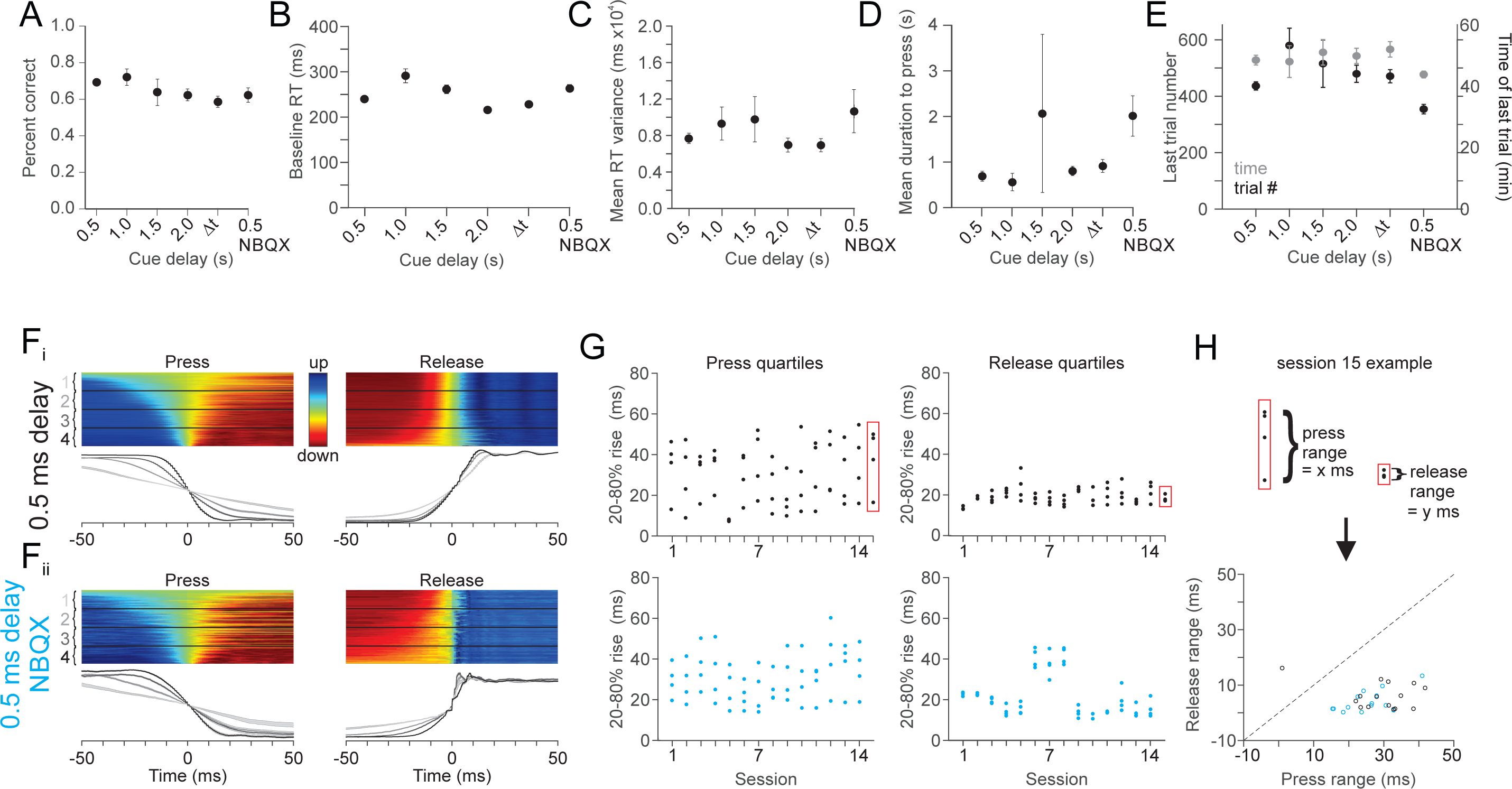
Performance in the cue prediction condition in control and NBQX sessions. A-E) Summary across experiments of mean percent correct lever releases, mean baseline reaction time from the visual cue, mean reaction time variance, mean duration to press, mean number of trials performed, and the mean duration of behavioral sessions. n=44 sessions, 0.5 s; n=9, 1.0 s; n=6, 1.5 s; n=30, 2.0 s; n=26, Δt; n=14, 0.5 s NBQX. Error bars are SEM across sessions. For each cue delay, the minimum and maximum (x,y) number of trials performed in single sessions was: 0.5 s: (309, 700) 1 s: (252, 900) 1.5 s: (202, 775) 2 s: (278,1185) Δt: (313, 772) 0.5 s NBQX: (227, 429). F_i_) Individual (top) and average (bottom) lever trajectories aligned to lever press (left) and release(right) segregated by duration into quartiles (n=112 trials / quartile) for a representative 0.5 s cue prediction session. F_ii_) Same as F_i_ for a representative 0.5 s cue prediction session in NBQX. (n=142 trials / quartile). G) Summary of press (left) and release (right) quartiles across experiments for control (top; n=15 sessions) and NBQX (bottom; n=14 sessions) experiments. H) Summary of the ratio between press and release ranges (difference between 1^st^ and 4^th^ quartile) for control (black) and NBQX (blue) experiments.

**Figure 4.**
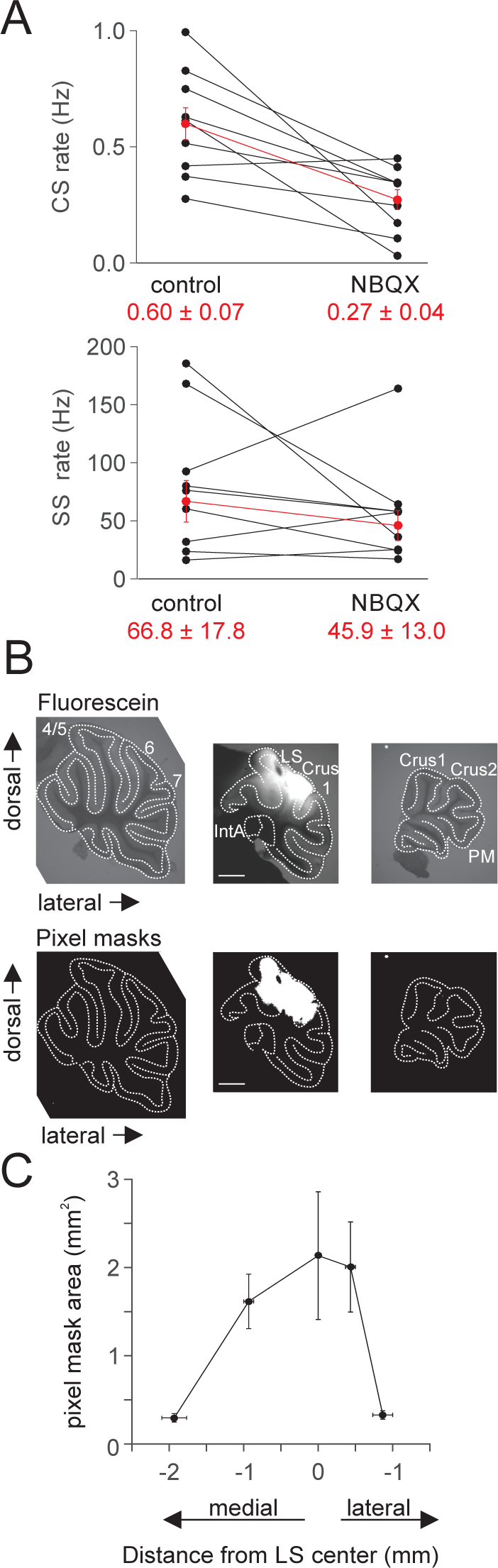
Pharmacological modulation of complex spiking in lobule simplex. A) Single unit recordings during NBQX application to LS. Top, summary of complex spike rates in control and after application of NBQX for individual cells (black; n=9 cells, 5 mice) and the average of all cells (red). Error bars are ±SEM across cells. Bottom, same as top for simple spike rates. Note that NBQX strongly reduces complex but not simple spike rates. B) (top) Raw epifluorescence images from parasagittal cerebellar sections showing fluorescein labeling after surface application to lobule simplex *in vivo* (methods, 3 mice). (bottom) Pixel masks from the same images above. Pixels were thresholded at 30% of maximum value to visualize and quantify fluorescein labeling. C) Summary of mediolateral fluorescein spread across sections. (n=3 animals). Error bars are SEM across animals.

**Figure 5.**
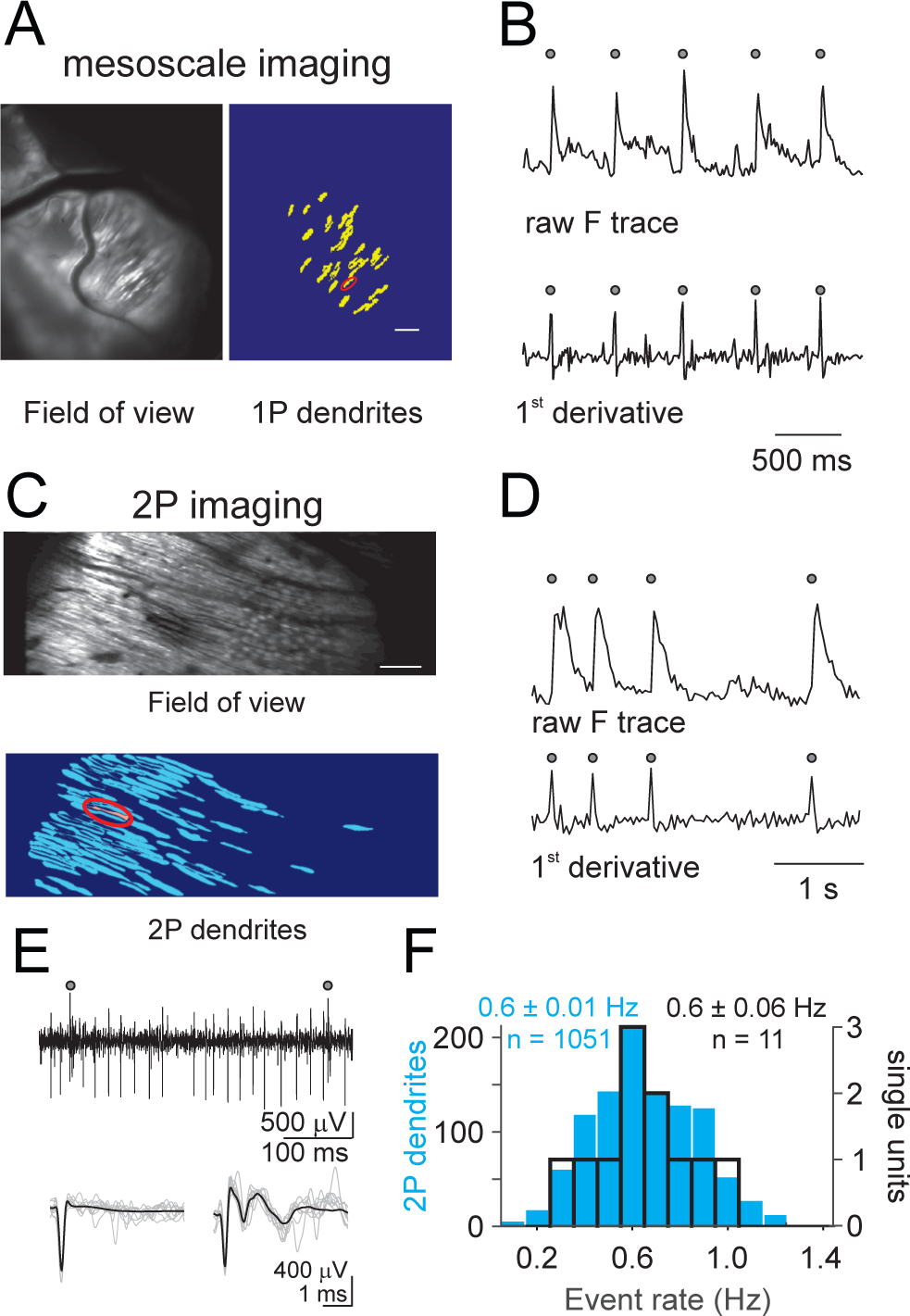
Segmentation of PC dendrites from imaging experiments. A) Example single photon imaging field of view (left) is segmented to mask individual dendrites (right) used for subsequent single cell analysis. Scale bar = 100 μm. B) Example timecourse of raw fluorescence (top) from the circled dendrite in A showing individual calcium transients identified (gray circles) according to peaks in the first derivative of the fluorescence trace (bottom). C) Same as A for an example two photon field of view. Note that while PC somata are visible (round, top right), image segmentation does not extract activity from these structures. Scale bar = 100 μm. D) Same as B for the experiment in C. E) Top, example time course of a single unit PC recording from an awake mouse illustrating detection of complex spikes (gray circles). Bottom, overlay of individual simple (left) and complex (right) spike waveforms (gray) and the average waveform (black). F) Summary of mean spike rates ± SEM across all two photon dendrite imaging sessions (blue; n=1146 dendrites) and acute single unit recordings (n=11 units).

**Figure 6.**
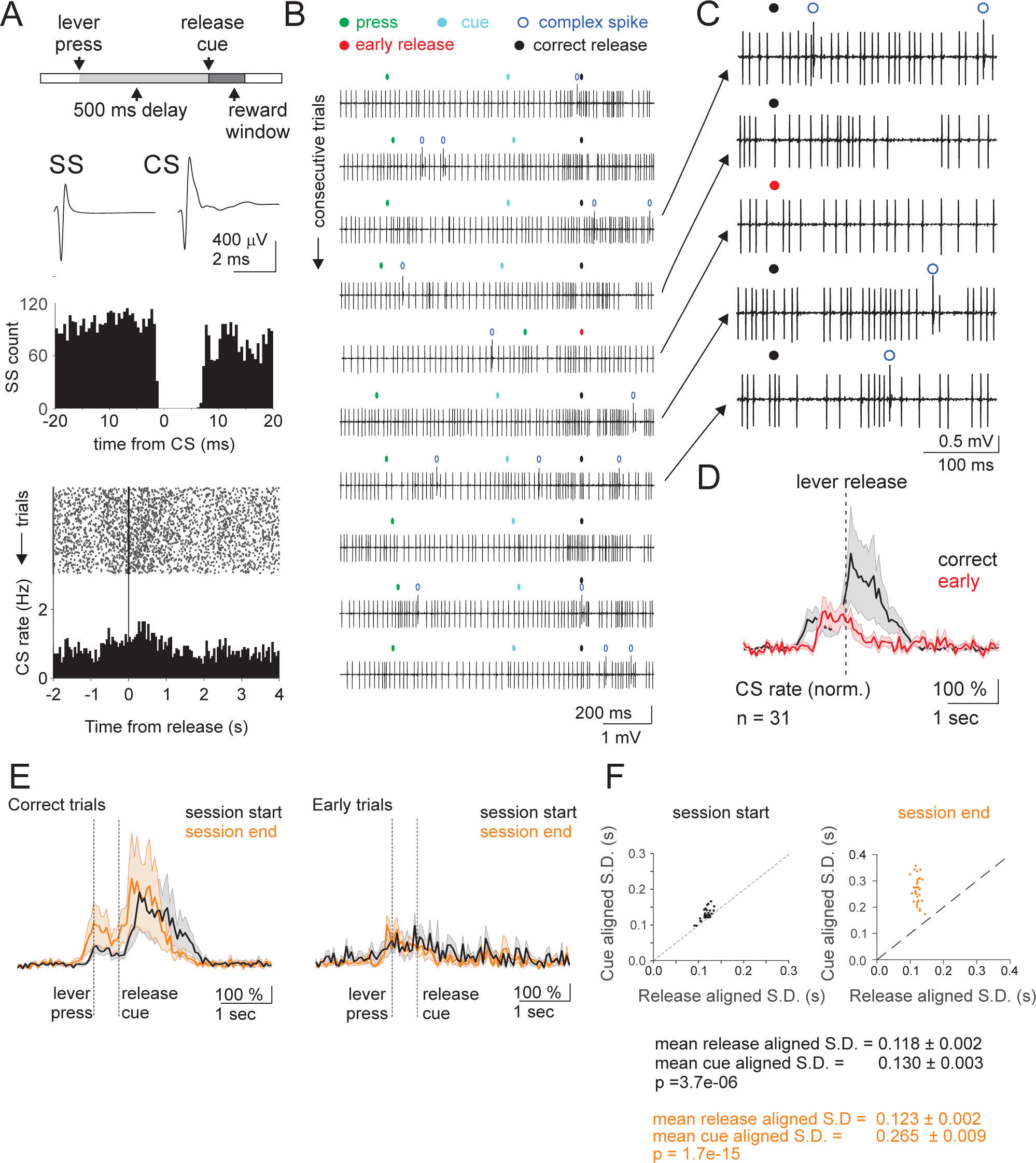
Single unit PC recordings of complex spiking during the cue prediction condition. A) Top, Schematic of the trial structure schematic, where a constant cue delay of 500ms was imposed on each trial. Middle top, Average simple (SS, left) and complex (CS, right) spike waveforms from an example single unit. Middle bottom, Histogram of SS firing rate aligned to complex spike time revealing post-CS pause, confirming isolation of single PC. Bottom, raster of single trial complex spikes and session PSTH aligned to lever release. B) Single trial voltage traces (complex spikes- open blue circle) from the cell in A aligned to lever release (correct- black circle; early- red circle). C) Expansion of 5 consecutive traces from B at time of lever release. D) Mean normalized complex spike rate (methods) from single unit recordings (31 PCs, 8 mice) aligned to lever release for correct (black) and early (red) trials during the cue prediction condition. Shaded error is SEM across PCs. E). Mean normalized complex spike rates aligned to cue presentation for the first 1/3 of trials (black) and the last 1/3 of trials (orange) across cue prediction sessions for correct (left) and early (right) trials (31 PCs, 8 mice). Shaded error is SEM across PCs. F) Summary of the S.D. of spike times when aligned to either lever release or visual cue for each PC (n=31). P-values reflect paired t-tests.

**Figure 7.**
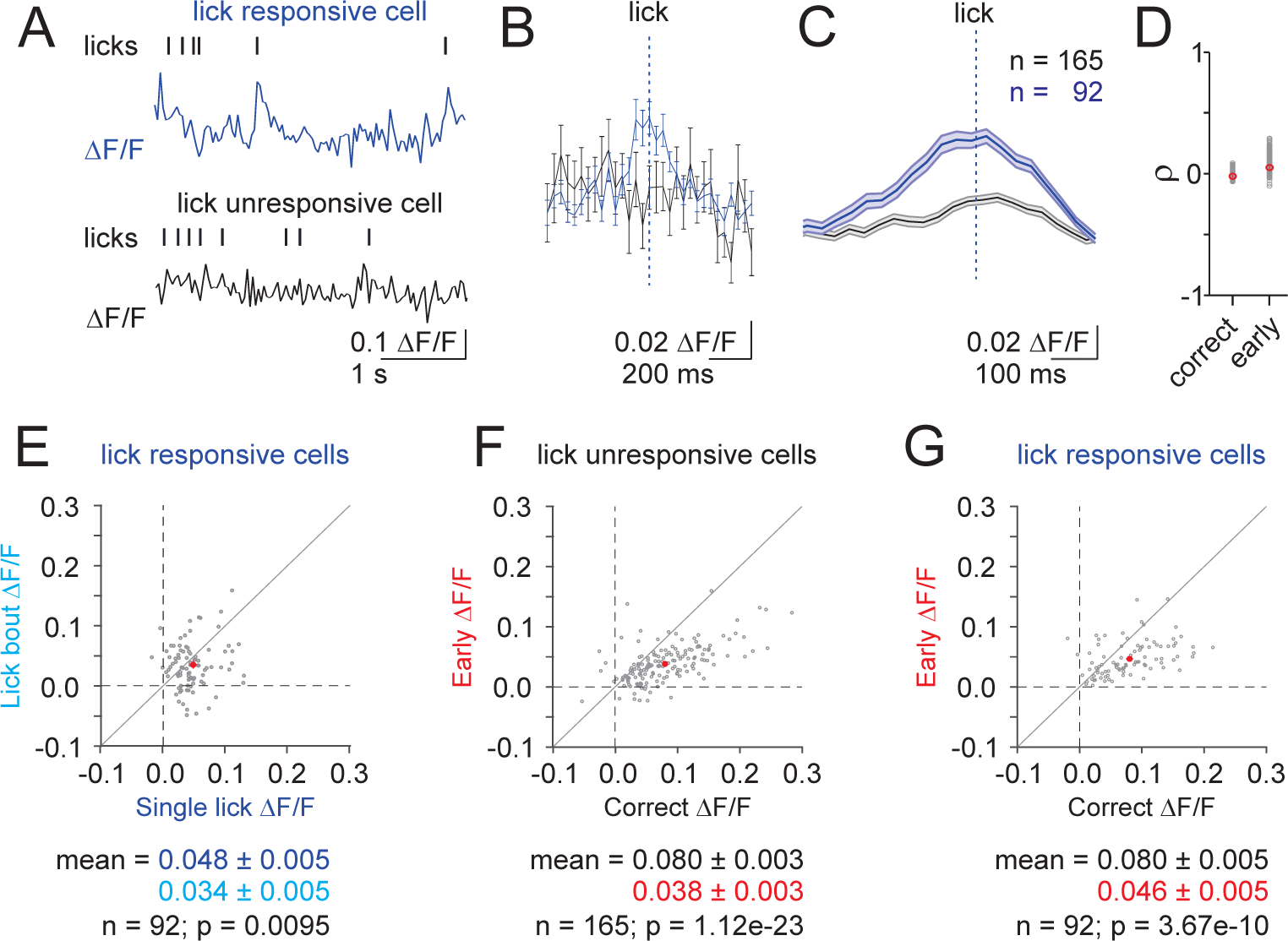
Differential complex spiking on correct and early release trials is not driven by motor signals due to licking. A) Example imaging data from an inter-trial interval (ITI) showing a lick responsive dendrite (top, dark blue) and a lick unresponsive dendrite (bottom, black) in the same field of view. B) Averaged lick triggered calcium transients for the lick responsive dendrite (dark blue) and a lick unresponsive dendrite (black) in A). n=28 licks, Error bars are SEM across lick-triggered events. C) Mean lick triggered calcium transient across all lick responsive (dark blue) and lick unresponsive (black) dendrites. Shaded error is SEM across dendrites. D) Summary of the Spearman’s correlation between lick rate and amplitude of the calcium transient in each frame across trials for lick responsive (dark blue) and lick unresponsive dendrites (grey). Red points indicate the mean ± SEM across dendrites (447 dendrites, 6 animals and 10 sessions). E) Summary comparing the amplitude of calcium transients for lick responsive neurons only in response to either a single lick (blue) or a lick bout (3 or more licks with ≤ 300 ms between licks; cyan). Note that additional licking does not produce larger responses. F) Summary comparing the amplitude of calcium transients on correct (black) and early (red) lever releases for lick unresponsive dendrites only. G) Same as F, for lick responsive dendrites. P-values reflect paired t-tests.

**Figure 8.**
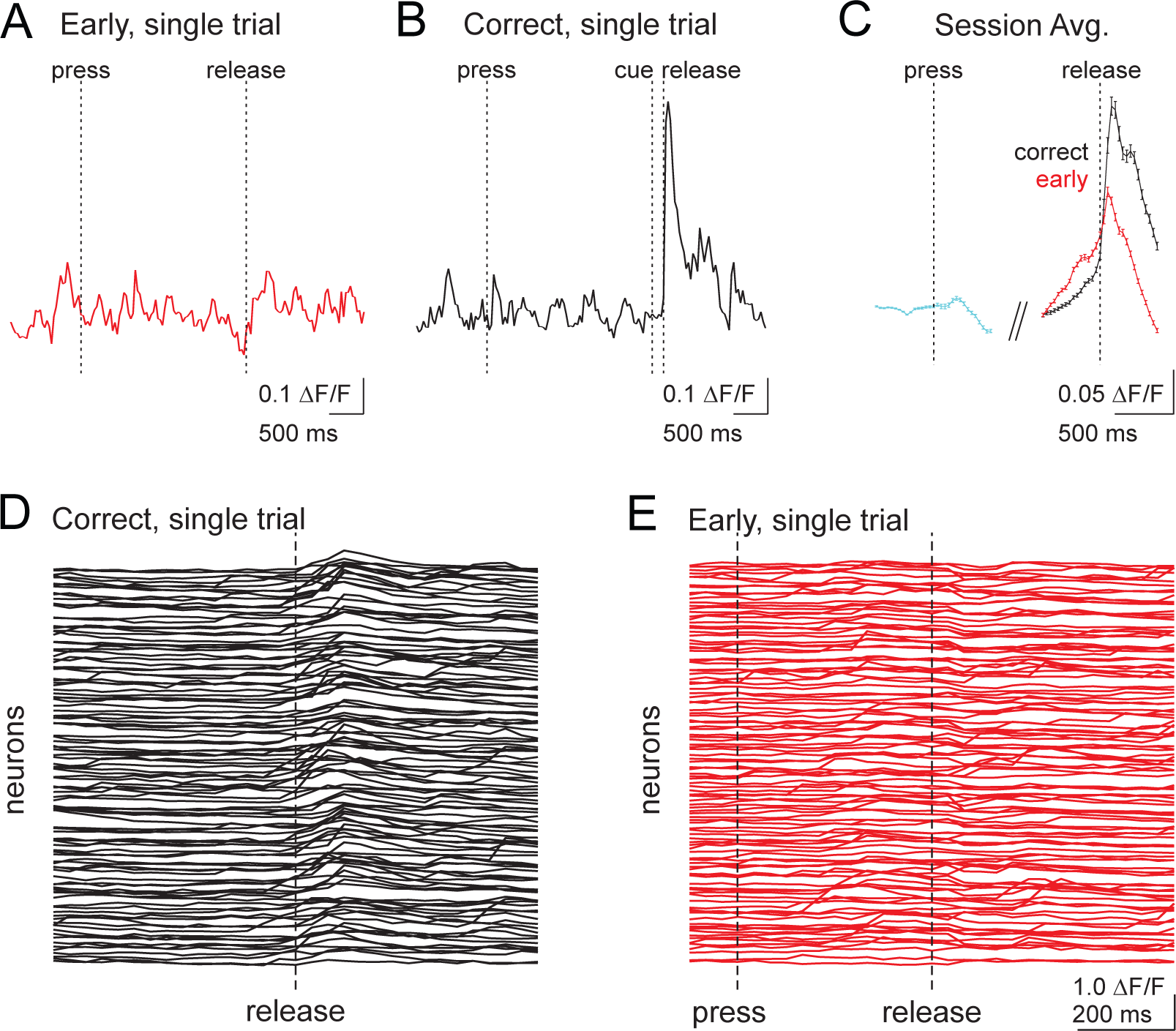
Example single trial and session average calcium transients from 2-photon imaging. A) Example single trial data from an individual dendrite showing the normalized calcium transient across an early release trial from before lever press to after the lever release. B) Same as A) but for a correct trial, including the time of cue presentation. C) Mean calcium transients across trials (n= 186) for the dendrite in A and B. Error bars are SEM across trials. D) Single trial example, aligned to lever release, showing all dendrites from the experiment illustrated in Fig. 5C on a correct trial over 1 second surrounding release (n=115 dendrites). Lever press is off scale. E) Same as D, but for an early release trial from the same imaging session.

**Figure 9.**
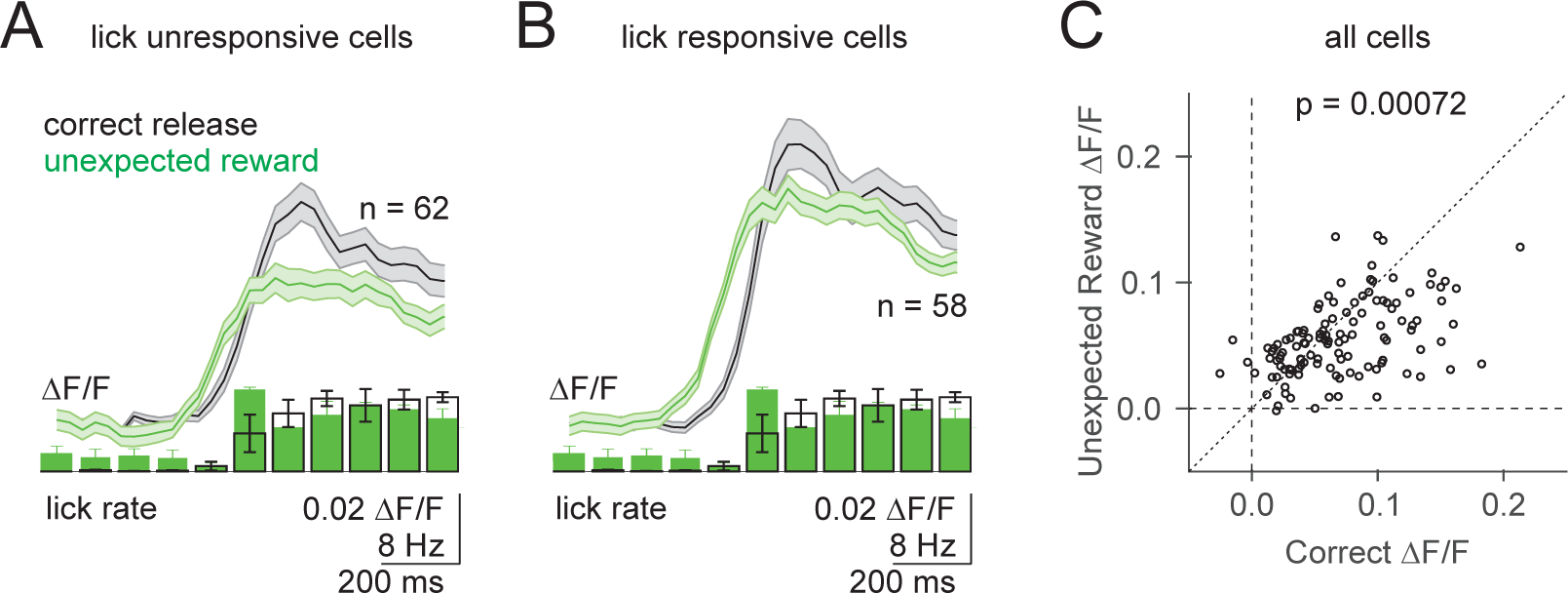
Unexpected reward drives complex spiking in lick unresponsive and responsive dendrites. A-B) Top, Average calcium transient in response to unexpected reward (green, aligned to first lick) and correct lever releases (black, aligned to release). Shaded error is SEM across dendrites. Bottom, Average lick rate for unexpected reward and correct lever releases. Error is SEM across experiments (n=3). Lick responsiveness was defined according to significant responses in the lick triggered averaged taken from the inter-trial interval (Supp. Fig. 7). C). Summary of peak calcium transients in the same neurons for correct lever releases vs unexpected reward trials. Note that responses are proportional, and response amplitude is determined by response probability (Fig. 5,6). P-value reflects paired t-test.

**Figure 10.**
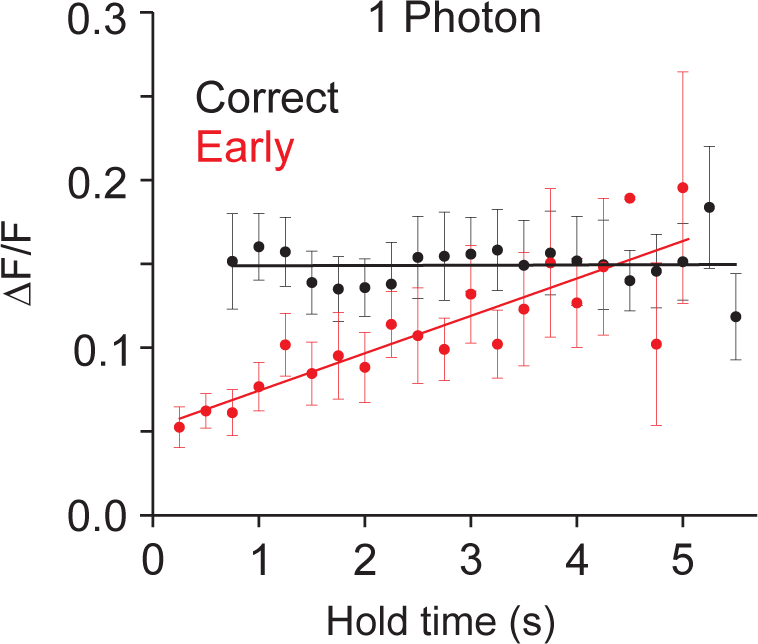
Complex spiking scales with lever hold time for early releases in single photon imaging experiments. Summary of peak calcium transients in a 500 ms window at the time of release (methods) across all single photon experiments for correct (black) and early (red) release trials binned according to hold time (250ms bins). Linear fits were applied to data from each trial type (n=10 animals, 17 sessions). Error bars are SEM across sessions.

**Figure 11.**
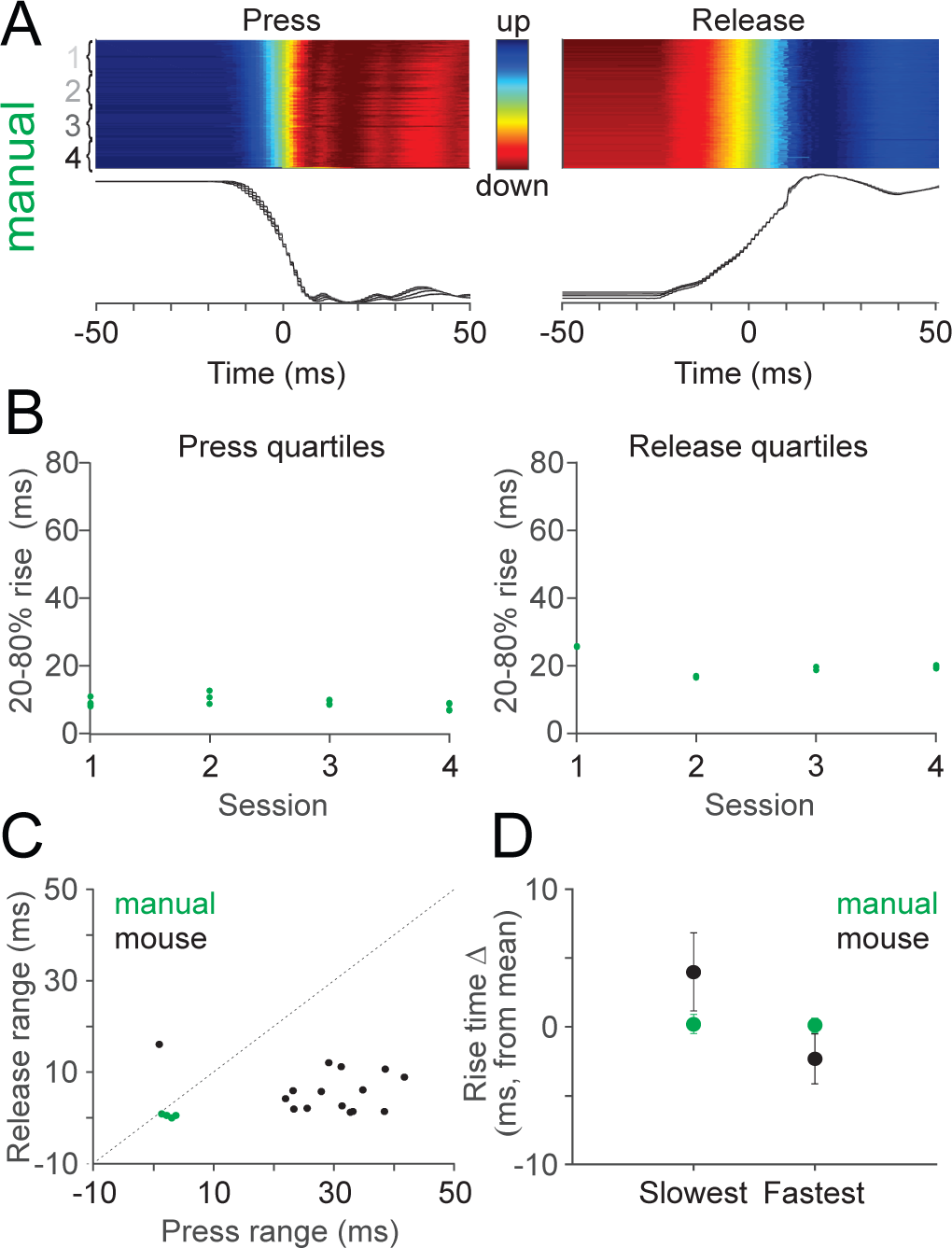
Mice have full control over the lever trajectory. A) Individual (top, n=217) and average (bottom, 54 per quartile) lever trajectories aligned to lever press (left) and release (right) for manual lever depression followed by unperturbed gravity return. Shaded error is SEM across trials. B) Summary of press (left) and release (right) quartiles across experiments for manual lever experiments (n=4). Note the lack of variability across quartiles. C) Summary scatterplot comparing 20-80% rise times of lever press and release for experiments where the lever was controlled by mice (black, n=15) and manual press (green, n=4) D) Summary comparison measuring the average difference from the mean of the slowest release quartile and the fastest release quartile for experiments where the lever was controlled by mice (black) and manual press (green). Lever trajectories were sorted according to duration in a window from 200ms before threshold crossing to the time of a threshold crossing half way between the top and bottom of the total lever displacement. The slowest and fastest 25 trials were extracted for comparison, and their average rise time was normalized by subtracting that of the mean trajectory of the whole session. Error bars are SEM across sessions. These results show that mice produce movements both slower and faster than the mean, indicating full control over the lever trajectory.

**Video 1.** Optogenetic silencing of superficial PCs in lobule simplex drives ipsilateral forelimb movements. An external fiber coupled laser was activated at the time indicated by the green circle above lobule simplex (methods), resulting in forelimb movement.

**Video 2.** Example correct and early release trials from the cue prediction condition. Two example trials from the same mouse in the same training session are shown first in real time, and then repeated at 0.25x video rate.

## Methods

### Mice

All experimental procedures using animals were carried out with the approval of the Duke University Animal Care and Use Committee. All experiments were performed during light cycle using adult mice (>p60) of both sexes, randomly selected from breeding litters. All mice were housed in a vivarium with normal light/dark cycles in cages with 1-5 mice. Imaging experiments were performed using Tg(PCP2-Cre)3555Jdhu mice (Jackson Labs, 010536; n=12). Optogenetic mapping experiments were conducted in PCP2-Cre animals crossed with Ai35(RCL-Arch/GFP) (Jax 012735; n=5 mice). Single unit recordings during the cue prediction condition were performed in wild type C57BL/6J mice (n= 8 mice). Additional behavioral experiments where imaging was not performed were conducted in wild type C57BL/6J mice (n=18). We used two exclusion criteria for animals in this study: (1) poor recovery or other health concerns following surgical intervention or (2) missed virus injection, as determined by *in vivo* imaging and post-hoc histological analysis.

### Surgical Procedures

3-10 hours prior to surgery, animals received dexamethasone (3mg/kg) and ketoprofen (5mg/kg). Surgical procedures were performed under anesthesia, using an initial dose of ketamine/xylazine (50mg/kg and 5mg/kg) 5 minutes prior to surgery and sustained during surgery with 1.0-2.0% isoflurane. Toe pinches and breathing were used to monitor anesthesia levels throughout surgeries. Body temperature was maintained using a heating pad (TC-1000 CWE Inc.). Custom-made titanium headplates (HE Parmer) were secured to the skull using Metabond (Parkell). For imaging experiments, a 3mm diameter craniotomy was made over the lobule simplex approximately 1.4mm lateral and 2.8mm posterior to lambda, and glass cover slips consisting of two 3mm bonded to a 5mm coverslip (Warner Instruments No. 1) with index matched adhesive (Norland No. 1) were secured in the craniotomy using Metabond. Buprenex (0.05mg/kg) and cefazolin (50mg/kg) were administered following surgery twice a day for two days. Following a minimum of 4 recovery days, animals were water deprived for 3 days, or until body weight stabilized at 85% of initial weight, and were habituated to head restraint (3-5 days) prior to behavioral training.

For imaging experiments, the glass cover slip was removed following behavioral training, and mice were injected (WPI UMP3) with AAV1.CAG.Flex.GCaMP6f.WPRE.SV40 (UPenn vector core, titer = 9.40×10^12^ or 7.60×10^12^). 150 nL virus diluted 1:1-1:5 in ACSF was injected at a rate of 30nl/min and a depth of 150 μm at 1-3 sites in dorsal lobule simplex. Imaging was performed beginning 14 days following injection.

For *in vivo* pharmacology experiments, a 3mm craniotomy was performed over lobule simplex of headposted mice, and the dura mater was peeled back at the center of the craniotomy. 10uL of combined NBQX (300 μM) and CPP (30 μM) was applied into a well surrounding the craniotomy on the surface of the cerebellum 20 minutes prior to behavioral training. In 8 of 14 experiments, MCPG (30 μM) was also included to block metabotropic glutamate receptors. Because there no significant differences in performance across these two groups, all data were pooled for analysis. During behavior, a second drug application of 10uL was administered 20 and 40 minutes into the task, and depending on the duration of training, a third dose was applied after 1 hour. Craniotomies were subsequently covered with silicone elastomer (WPI, Inc.) prior to returning animals to their home cage. To quantify the spread of pharmacological agents in the cerebellum, 10 μL of fluorescein dye (Sigma-Aldrich #F6377, 1mM in aCSF) was applied to the craniotomy in 3 animals with the same method and timecourse as for drug applications. Following standard histological processing and imaging (below), labeling was quantified by creating a binary pixel mask thresholded at 30% of the maximum fluorescence value for each experiment. Dye labeling was then measured using the area of these pixel masks (Supp. Fig. 4 B,C).

### Behavior

During behavioral training, animals were head-fixed and placed in front of a computer monitor, lever and reward delivery tube. Animals were trained to self-initiate trials by depressing the lever using their right forepaw, and required to successfully hold the lever in the down position for randomized intervals ranging between 500ms and 5 s on the cue reaction paradigm before performing the cue prediction paradigm. For both paradigms, a high contrast hold cue was present at all times, including the intertrial interval (ITI), and transitioned 90 degrees to the release cue on each trial at the instructed time of release until the animal either released the lever or 1 second had passed. Training sessions lasted for 90 minutes, and learning in the cue prediction task occurred with the range of 309-700 trials. Lever releases within one second of the release cue were rewarded immediately at the time of lever release (0.01 M saccharine). Immediately following any lever release, a solenoid was engaged to prevent lever press during the ITI. Following the ITI (3-6 s), the solenoid is lowered, allowing the mouse to self-initiate a new trial. During training, a 1-3 second ‘timeout’ was implemented to punish early lever releases. No timeouts were used in fully trained animals for imaging or behavior data collection sessions. For reward omission sessions, 20% of randomly determined correct lever releases were unrewarded. Animals used for imaging experiments performed the cue reaction condition with a mean peak percent correct of 79.7 ± 0.23% achieved in 26 ± 2 training days. On imaging days animals performed a range of 150 to 580 trials per session with a mean of 273.9 ±13.1 trials. Behavioral parameters including lever press, lever release and licking were monitored using Mworks (http://mworks-project.org) and custom software written in MATLAB (Mathworks). To assess the degree of lever control by the mice, the dynamics of lever press and release trajectories were compared to lever presses initiated by the experimenter where the lever was allowed to return to the rest position on its own (Supp. Fig. 11). These results demonstrate that mice moved the lever both faster and slower than its intrinsic kinematics, and thus had full control over its trajectory. Licking was measured with electrical contact circuit. Inter-trial intervals ranged from 3 to 5 seconds.

### Calcium Imaging

#### Wide-field imaging

Single photon imaging was performed using a customized microscope (Sutter SOM) affixed with a 5x objective (Mitutoyo, 0.14NA) and CMOS camera (Qimaging, Rolera em-c^2^). Excitation (470 nm) was provided by an LED (ThorLabs, M470L3), and data were collected through a green filter (520-536 nm band pass, Edmund Optics) at a frame rate of 10Hz, with a field of view of 3.5×3.5mm at 1002×1004 pixels.

#### Two-Photon Imaging

Two-photon imaging was performed with a resonant scanning microscope (Neurolabware) using a 16x water immersion objective (Nikon CFI75 LWD 16xW 0.80NA). Imaging was performed using a polymer to stabilize the immersion solution (MakingCosmetics, 0.4% Carbomer 940). A Ti:Sapphire laser tuned to 920nm (Spectra Physics, Mai Tai eHP DeepSee) was raster scanned via a resonant galvanometer (8 kHz, Cambridge Technology) onto the brain at a frame rate of either 30 Hz with a field of view of either 1030 μm x 581 μm (796 × 264 pixels) or 555 μm x 233 μm (796 × 264 pixels), or 15.5Hz with a field of view of 555 μm x 452 μm and (796 × 512 pixels). Data were collected through a green filter (510 ± 42 nm band filter (Semrock)) onto GaAsP photomultipliers (H10770B-40, Hamamatsu) using Scanbox software (Neurolabware). A total of 12 mice were used for imaging experiments (10 mice for wide-field, 11 mice for two-photon, 12 mice total).

### Single Unit Recordings and Optogenetics

Acute single unit recordings were performed in awake animals by performing a craniotomy over lobule simplex and inserting a multi-electrode silicone probe (Neuronexus, A4×8-5mm-100-400- 177-A32, 4 shanks, 8 site/shank at 100 μm spacing) using a Cerebus multichannel acquisition system (Blackrock Microsystems, Salt Lake City). For single unit recordings obtained in the cue prediction condition (Supp. Fig. 6), chronically implanted electrode arrays were used (Dual drive movable electrode bundles with 8 tungsten electrodes (23 μm) in each cannula or 16 tungsten electrodes bundle in one cannula, Innovative Neurophysioloy Inc). Electrode arrays were implanted using stereotaxic coordinates to target lobule simplex at AP 6.2; ML 2.0. Electrode bundles were inserted into lobule simplex at a depth of approximately 0.2-0.3 mm. The implant was encased in Metabond for stability.

For both recording conditions, continuous recording data was bandpass filtered with a 2- pole Butterworth between 250 Hz and 5 kHz and referenced against an electrode with no spikes using Spike2. Single units were isolated by amplitude thresholding. Template waveforms were defined and characterized by their width and peak, and PCA of waveforms was done in off-line in Spike2.

Complex spikes and simple spikes were discriminated as in de Solages et al.^50^, first on the basis of their stereotypical waveform. Complex spikes typically had a multi-wavelet form, including a large positive peak within 6ms following spike initiation. Manual identification of 10- 20 complex spikes was used to generate a mean template waveform, which was then compared to all other spikes for a given unit using the Spearman rank order correlation coefficient. The combination of Spearman coefficients and the magnitude of the positive waveform deflection was used to segregate complex and simple spikes. The presence of a post-complex spike pause (20 ms or more) in simple spike firing was verified by cross-correlogram for all isolated single units.

Complex spike rates for the cue prediction condition were normalized to the baseline firing rate determined in a one-second window during the ITI according to: (FR-baseline)/baseline) (Supp. Fig. 6).

For optogenetic mapping of the dorsal cerebellum, an optical fiber (0.39 NA, 400 µm core multimode, ThorLabs FT400EMT) coupled to a 532 nm laser (Optoengine, MGL-III-532) was positioned above the cranial window using a micromanipulator (Scientifica PatchStar).

### Histology

Mice were deeply anesthetized with ketamine/xylazine (200mg/kg & 30mg/kg respectively, IP) then perfused with PBS then 4% paraformaldehyde. 100um sagittal sections were cut using a vibratome (Pelco 102). Cerebellar slices were then mounted using a mounting medium (Southern Biotech Fluoromount-G or DAPI Fluoromount-G) and imaged with a fluorescence microscope (Nikon Eclipse 80i).

### Data Analysis and Statistics

#### Behavior

Behavior sessions were only analyzed within the time range of active task performance. Accordingly, the last trial of each session was determined by the occurrence of either two consecutive failed trials (in which the mouse did not release the lever in response to the cue) or two consecutive trials with post-ITI duration to press longer than 1.5x the session average.

Reaction times were measured using lever releases within the reward window as well as those occurring 200ms prior to the reward cue to account for predictive responses. Reaction times are plotted as a binned average of five trials. Initial reaction times were calculated according to the y-intercept of the linear regression of the first fifty trials. Final reaction time at the end of each session was calculated by averaging the reaction times of trials 200-250.

The symmetry of reaction time data about the sample mean was measured to test the extinguishment of learning after following cue prediction sessions according to skewness. Sample skewness was defined by

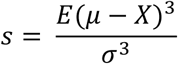

where *µ* is the mean of *X, E*(∗) is the expected value of the quantity ∗, and *σ* is the standard deviation of *X*. Thus, for a given sample *x*, this calculates:

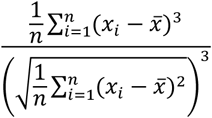

for *x*_*i*_ ∈ *x*, where 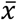 is the sample mean.

Movement trajectories were measured using a lever affixed to a rotary encoder (US Digital) with a 60mm radius and 1250 pulses per revolution. Encoder values were collected at 1000 Hz, and trajectories were calculated by up-sampling at 200 μs (5000 Hz) using nearest-neighbor interpolation. Pulse number was converted to degrees, and the vertical displacement in millimeters was calculated using the chord length of the leaver displacement angle. 20-80% rise times were calculated using normalized average trajectories. Presses and releases were sorted according to duration in a window from 200ms before threshold crossing to the time of a threshold crossing half way between the top and bottom of the total lever displacement.

#### Wide-field imaging

Imaging analysis were performed using custom MATLAB code. Regions of interest (ROIs) were selected within empirically defined forelimb movement region of lobule simplex according to the spread of GCaMP expression. Window location and the lobule identity were identified according to folia patterns visible through the cranial window, landmarks recorded during surgery, and post-hoc histology. Baseline fluorescence (F) was measured on a trial-by-trial basis during the inter-trial interval (ITI) as the mean fluorescence 900 to 200ms before trial initiation. Normalized fluorescence (ΔF/F) was calculated according to the cumulative activity within an ROI, or on a pixel by pixel basis. Lever hold times <200ms in total, or > 1 s after the visual cue were exceedingly rare, and thus excluded from analysis. Lever releases between 200ms and 1000ms following the visual cue were classified as correct, and releases prior to visual cue were classified as early. Lever releases ≤200 ms following the visual cue were considered too fast to be reactions to the cue based on average reaction time distributions. Peak ΔF/F was measured on a trial by trial basis in the time window from 100ms before to 400 ms after lever release. Note that differences in the timing of single trial calcium transient peaks produce smaller amplitudes for ΔF/F timecourses as compared to reported peak ΔF/F measurements. Spearman’s correlation between lick rate and peak ΔF/F within sessions was calculated according to the lick rate within 500 ms following each lever release and the peak calcium transient for each trial. For this correlation analysis, trials without licking were removed, and only sessions with 7 or more trials in each condition were included.

#### Meta K-means analysis and Cluster Correlations

Images were first registered to reduce motion artifacts, and then thresholded at 70% of maximum intensity for each frame to remove background noise. Images were downsampled 5-fold in both X and Y. Baseline F was defined as the averaged fluorescence across the entire movie for each pixel, and used to normalize change in fluorescence (ΔF/F). ΔF/F was then re-normalized to the maximum ΔF/F during the entire movie for each pixel. The repeated k-means clustering algorithm (meta-k-means) separated pixels to cluster centroids based on a pairwise correlation distance function using Pearson’s linear correlation coefficient (r). The final clusters were determined by first thresholding all the k-means results at 800 out of 1000 runs, and then by merging highly correlated clusters based on Dunn’s index. Clusters occupying less than 3% of the total imaging field were excluded from further analysis. Intra-cluster and inter-cluster correlation coefficients were calculated between 100 ms before and 300 ms after lever release for both correct and early trials on a frame by frame basis. To test for differences in the relationship between spike rates and correlation coefficients between trial types, paired clusters from each distribution were resampled with replacement and fit with a line to measure the y-intercept 1000 times. Statistical significance was computed according to the 95% confidence interval of the distribution of y-intercept difference values (difference between y-intercepts 0.0375 ± 0.0013, 95% confidence interval [0.0349, 0.0402]).

#### Two-photon imaging

Motion in the XY plane was corrected by sub-pixel image registration. To isolate signals from individual PC dendrites, we utilized principal component analysis (PCA) followed by independent component analysis (ICA). Final dendrite segmentation was achieved by thresholding the smoothed spatial filters from ICA. A binary mask was created by combining highly correlated pixels (correlation coefficient > 0.8) and removing any overlapped regions between segmented dendrites. Notably, image segmentation using these criteria did not extract PC soma, which were clearly visible in many single- and two-photon imaging experiments. Fluorescence changes (ΔF) were normalized to a window of baseline fluorescence (F) between 500ms and 100ms preceding trial initiation. Responses were categorized as significant (p<0.05) according to a one-tailed t-test if the ΔF/F in a 200 ms window surrounding lever press or release was larger than that from a 500 ms window immediately preceding lever movement. To extract events on single trials used for estimation of complex spike rates and event amplitudes, the first derivative of the raw fluorescence trace was thresholded at 2.5 standard deviations from baseline. The same methods were used to extract events for the subset of single photon experiments described in Supp. Fig. 5. For single trial amplitude measurements, only events well separated in time (greater than 650ms from the next event) were considered; however, for measurements of rates, all events separated by at least one frame were included. Note that rate estimates are thus likely to be lower than actual rates, particularly for single photon imaging where data were collected at 100 ms intervals. For measurements of standard deviation of event times, events were extracted from a window 433 ms around the lever release, and independently aligned to the time of visual cue. The same criteria were applied to electrophysiological measurements of spike times. Event latencies were calculated according to the time of peak event probability. Peak calcium transients on reward omission trials and on early release trials were quantified in two time windows (Fig. 8). Window 1 spanned the first 100 ms after lever release, and window 2 spanned 175 ms that began 165ms after lever release. Spearman’s correlation was calculated between lick rate and the peak calcium transient within sessions on a frame by frame basis for each trial in a window spanning 500 ms after lever release. Correlation values for correct and early release trials for each dendrite were averaged separately. Some neurons showed responses to licking, as defined by a mean lick- triggered calcium transient (for dendrites with at least 4 lick events per session) that was significantly larger (p<0.05) with respect to baseline (Supp. Fig. 7). Lick triggered averages were constructed from licks presumed to be spontaneous that occurred during the inter-trial interval with a buffer of 1000 ms between the end of the previous trial and the start of analysis to avoid contamination of reward-related licking.

#### Additional statistics

Data are presented as mean ± S.E.M., unless stated otherwise. Statistical tests were two-sided, except as specifically noted, and analyses of variance (ANOVA) were performed when more than two groups were tested. Differences were considered statistically significant when *P* < 0.05. No correction for multiple comparisons was applied. No statistical methods were used to predetermine sample sizes. Data distribution was assumed to be normal but this was not formally tested. Data collection and analysis were not performed blind to the conditions of the experiments, but data collection relied on automatized measurements and subsequent analysis was based on code uniformly applied across experimental conditions.

